# Bruce suppresses autophagy-regulated caspase activity and wing tissue growth in *Drosophila*

**DOI:** 10.1101/2025.08.24.672027

**Authors:** Natsuki Shinoda, Yutaro Hama, Nozomi Hanawa, Masayuki Miura

## Abstract

Caspases are cysteine-aspartic proteases that mediate both lethal and non-lethal cellular outcomes, including the promotion of tissue growth. However, the mechanisms underlying the differential regulation of these activities remain unclear. We have previously shown that among the two *Drosophila* executioner caspases, Dcp-1 and Drice, Dcp-1 promotes tissue growth in a non-lethal manner, independent of canonical apoptotic signaling. Herein, we demonstrated that overexpressed Dcp-1, but not Drice, was activated without canonical apoptosome components. TurboID-based proximity labeling revealed distinct proximal proteomes, among which Sirtuin 1, an Atg8a deacetylase, which promotes autophagy, was specifically required for Dcp-1 activation. Autophagy-related genes, including Bcl-2 family members *Debcl* and *Buffy*, are required for Dcp-1 activation. Structure-based prediction using AlphaFold3 further identified Bruce, an autophagy-regulated inhibitor of apoptosis, as a Dcp-1-specific regulator acting outside the apoptosome-mediated pathway. Physiologically, Bruce suppresses wing tissue growth. These findings indicate that non-lethal Dcp-1 activity is governed by the autophagy-Bruce axis, enabling distinct non-lethal functions independent of cell death.

## Introduction

Caspases are cysteine-aspartic acid proteases that serve as the central executioners of apoptosis. In *Drosophila*, core caspase-mediated signaling for apoptosis is well conserved (Nakajima and Kuranaga, 2017; Shinoda and Miura, 2024). Upon apoptotic stimulation, the pro-apoptotic genes *rpr*, *hid*, and *grim* (*RHG*), which antagonize DIAP-1, an inhibitor of apoptosis protein (IAP), are transcriptionally upregulated. The degradation of DIAP-1 allows the initiator caspase Dronc and its activator Dark to assemble into the apoptosome, thereby triggering a proteolytic cascade that activates the executioner caspases Dcp-1 and Drice, ultimately leading to apoptosis. In addition to their role in apoptosis, caspases perform diverse non-cell death functions collectively termed caspase-dependent non-lethal cellular processes (Aram et al., 2017). However, the mechanisms that differentially regulate lethal and non-lethal caspase activity remain largely unknown.

Functional specificity appears to arise at the level of the executioner caspases. Of the two executioner caspases in *Drosophila*, Drice is more potent than Dcp-1 in inducing cell death in wing imaginal discs (Florentin and Arama, 2012). In contrast, Dcp-1 is required for autophagy triggered by starvation or proteasome dysfunction in the ovaries (Choutka et al., 2017; DeVorkin et al., 2014; Hou et al., 2008). We have previously demonstrated that Dcp-1 and another executioner caspase, Decay, but not Drice, specifically promote imaginal tissue growth independent of apoptosome and cell death (Shinoda et al., 2019). Mechanistically, we showed that cleavage-mediated inactivation of Acinus, a positive regulator of autophagy (Haberman et al., 2010; Nandi et al., 2014), which limits tissue growth independently of autophagy (Tyra et al., 2020), partly promotes tissue growth (Shinoda et al., 2019). However, the upstream regulatory mechanisms that selectively regulate Dcp-1 and Decay activity remain unknown.

Proximity labeling has emerged as a powerful approach for comprehensively analyzing molecular interactions, including protein-protein interactions (Qin et al., 2021). TurboID enables the rapid biotinylation of lysine residues on neighboring proteins within 10 minutes in cultured cells (Branon et al., 2018). Streptavidin-based enrichment of these biotinylated proteins, followed by mass spectrometry (MS), allows the proteome-wide identification of proteins near a target of interest. We previously generated *Drosophila* lines in which TurboID was knocked in at the C-termini of Dronc, Dcp-1, or Drice (Shinoda et al., 2019). In wing imaginal discs, TurboID-based labeling has revealed distinct proximity patterns for Dcp-1 and Drice, suggesting that each executioner caspase is associated with a unique set of neighboring proteins (Shinoda et al., 2019). We have shown that the differences in substrate specificity between Dcp-1 and Drice were consistent with their distinct proximity profiles in the ovary, as exemplified by BubR1 (Shinoda et al., 2023). Moreover, overexpression of *Fasciclin 3*, a Drice-proximal protein identified in adult brains, induces non-lethal caspase activation in a Dronc-dependent manner (Muramoto et al., 2025). These findings indicate that caspase-proximal proteins can function not only as potential substrates but also as regulatory factors.

In this study, to dissect how executioner caspase activity is differentially regulated to support non-lethal functions versus apoptotic cell death, we focused on the two executioner caspases that are known to participate in apoptosis, Dcp-1 and Drice. More specifically, we investigated the regulatory mechanisms underlying Dcp-1 activation. We found that the overexpression of *Dcp-1* induced a wingless phenotype. Using this phenotype as a functional readout, we found that Dcp-1 activation specifically requires its proximal protein Sirtuin 1 (Sirt1), Bcl-2 family members Debcl and Buffy, and components of the autophagy pathway. Notably, the autophagy-regulated giant IAP Bruce specifically interacts with and regulates Dcp-1 activity without affecting apoptosome-mediated core apoptotic signaling, thereby suppressing wing imaginal tissue growth. Together, these findings reveal that Bruce acts upstream to diverge the regulatory pathways of Dcp-1 and Drice, enabling Dcp-1 to mediate distinct non-lethal functions through autophagy, independent of cell death.

## Results

### Overexpression of Dcp-1, but not Drice, induces apoptosis

Structurally, the executioner caspases, Dcp-1 and Drice, are similar, each comprising catalytic P20 and P10 domains and a highly disordered N-terminal prodomain (Figure 1A). First, we assessed the ability of two executioner caspases to trigger apoptosis. Using the *WP-Gal4* driver, which is expressed in the wing pouch region of the wing imaginal disc (Obata et al., 2014), overexpression of *Dcp-1* resulted in a wingless phenotype whereas the overexpression of *Drice* did not (Figure 1B). In wing imaginal discs, *Dcp-1* overexpression induces nuclear condensation (Figure 1C), indicative of apoptotic cell death. To further characterize the type of cell death, we examined whether overexpressed Dcp-1 is activated through cleavage. By western blotting, we found that Dcp-1 is cleaved upon overexpression (Figure 1–figure supplement 1A). We also found that simultaneous overexpression of the caspase inhibitor *p35* suppressed Dcp-1 cleavage (Figure 1–figure supplement 1A), suggesting that Dcp-1 is cleaved by caspases. Cleavage of overexpressed Dcp-1 is further confirmed by anti-cleaved Dcp-1 antibody staining in wing imaginal discs (Figure 1–figure supplement 1B), indicating that overexpressed Dcp-1 is cleaved and thereby activated. To evaluate whether overexpressed Dcp-1 shows typical executioner caspase activity, we employed two executioner caspase activity probes: GC3Ai (Schott et al., 2017; Zhang et al., 2013) and CD8::PARP::VENUS (Williams et al., 2006). GC3Ai becomes fluorescent upon caspase-mediated cleavage (Figure 1–figure supplement 1C). In wing imaginal discs, *Dcp-1* overexpression resulted in strong GC3Ai fluorescence, suggesting that it exerts typical executioner caspase activity (Figure 1–figure supplement 1D). CD8::PARP::VENUS generates a neo-epitope that can be detected by anti-cleaved PARP antibody upon caspase-mediated cleavage (Figure 1–figure supplement 1E). Consistently, in wing imaginal discs, *Dcp-1* overexpression led to a strong anti-cleaved PARP signal, further supporting that it induces typical executioner caspase activity. Consistent with these results, Dcp-1 overexpression resulted in TUNEL-positive cell death, reflecting DNA strand breaks that are a hallmark of apoptosis (Figure 1D). Overall, these results suggest that *Dcp-1* overexpression is sufficient to induce typical executioner caspase activity-dependent apoptotic cell death, consistent with our previous observations in olfactory receptor neurons (Muramoto et al., 2025).

**Figure 1.**
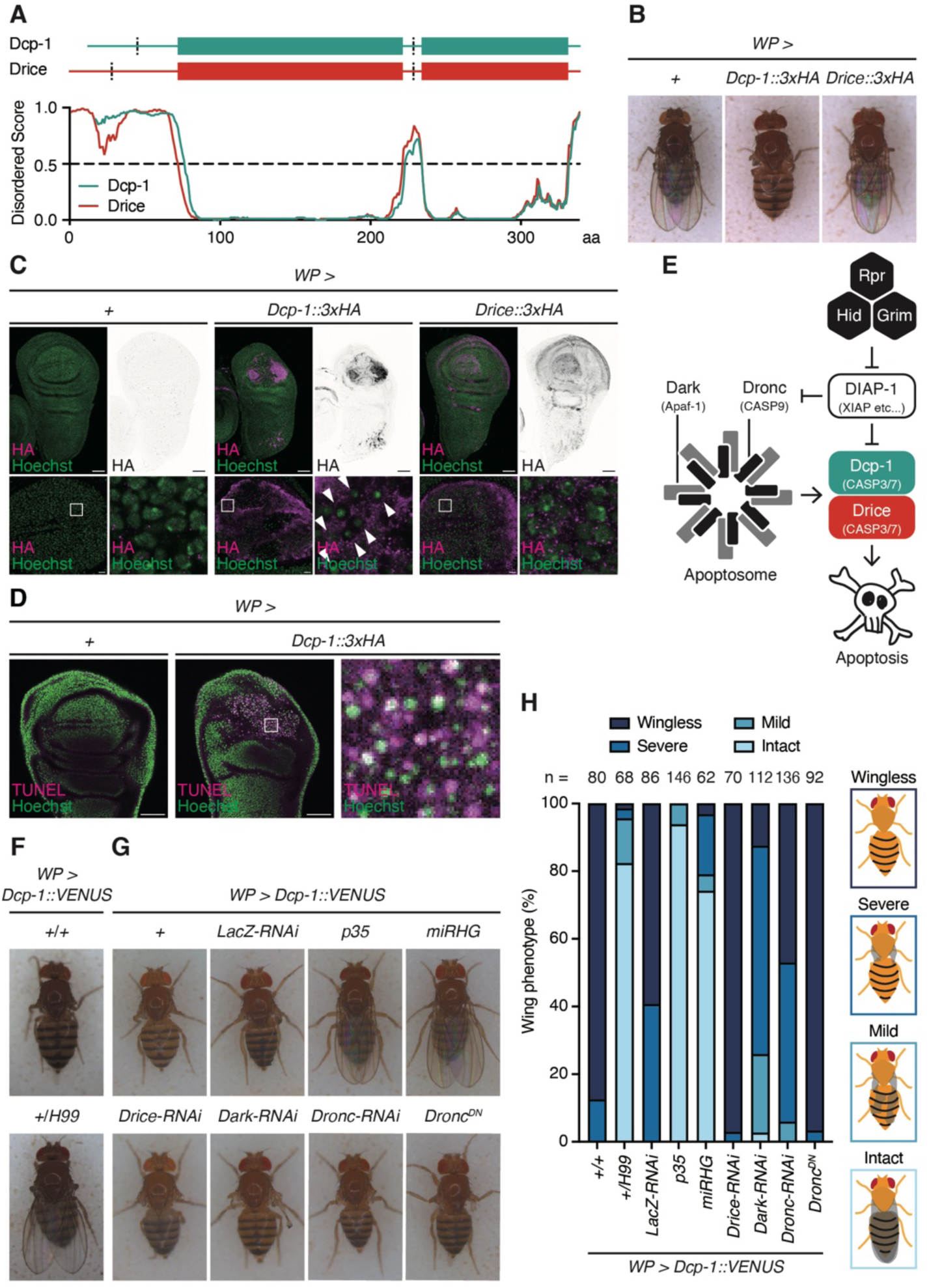
*Dcp-1* overexpression induces apoptosis independent of the apoptosome. (A) Schematic structures and intrinsically disordered region prediction of Dcp-1 (dark cyan) and Drice (red). The predicted disordered score plot is shown. Regions wherein the score is more than 0.5 are predicted to be disordered. (B) Representative images of female wing phenotypes upon caspase overexpression using *WP-Gal4*. (C) Representative images of larval wing imaginal disc upon *3xHA-tagged executioner caspase* (magenta) overexpression using *WP-Gal4*. Nuclei are visualized by Hoechst 33342 (green). Magnified images are shown in right below. Arrowheads indicate condensed nuclei. Scale bar: 50 µm (top) and 10 µm (bottom). (D) Representative images of the TUNEL staining (magenta) of larval wing imaginal discs upon *3xHA-tagged Dcp-1* overexpression using *WP-Gal4*. Nuclei are visualized by Hoechst 33342 (green). Magnified images are shown in right. Scale bar: 50 µm. (E) Schematic diagram of canonical apoptosis signaling in *Drosophila*. (F, G) Representative images of female wing phenotypes upon *Dcp-1::VENUS* overexpression using *WP-Gal4* with the genetic perturbation of canonical apoptosis signaling. (H) Quantification of (F) and (G). Each wing is manually classified into four (wingless, severe, mild, and intact) categories. Schematic images of the four categories are shown on the right. Sample sizes are shown in the figure.

Next, we examined the apoptotic potential of fluorescently tagged caspases in *Drosophila* S2 cells expressing mNeonGreen (mNG)-fused Dcp-1 or Drice. Dcp-1 expression caused a marked reduction in mNG intensity (Figure 1–figure supplement 2A–C) and a decreased number of mNG-positive cells (Figure 1–figure supplement 2A, D). These effects required Dcp-1 catalytic activity, as the mutation of the catalytic cysteine to glycine restored both mNG intensity and cell number (Figure 1–figure supplement 2A–D). In contrast, Drice expression neither reduced mNG intensity nor decreased the number of mNG-positive cells, resembling the catalytically inactive mutant (Figure 1–figure supplement 2A–D). Taken together, these results demonstrate that Dcp-1, but not Drice, is activated upon overexpression and induces apoptosis in multiple cell types.

### Dcp-1 activation is independent of the apoptosome composed of Dronc and Dark

Executioner caspases are typically activated by cleavage mediated by the initiator caspase, Dronc, which is activated through the assembly of the Dark-dependent apoptosome (Figure 1E) (Nakajima and Kuranaga, 2017; Shinoda and Miura, 2024). We previously proposed that Dcp-1 can be activated independently of Dark and Dronc (Shinoda et al., 2019). To test this hypothesis further, we analyzed genetic modifiers of the Dcp-1-induced wingless phenotype. A heterozygous mutation in the *H99* deficiency, which encompasses the pro-apoptotic *reape*r, *hid*, and *grim* (*RHG*) genes, rescued the wingless phenotype caused by *Dcp-1* overexpression (Figure 1F, H). Similarly, expression of the caspase inhibitor *p35* or knockdown of *RHG* genes also rescued this phenotype (Figure 1G, H), suggesting that Dcp-1 induces apoptosis in a catalytic activity-dependent manner under the control of RHG proteins. In contrast, knockdown of other core apoptotic components, including *Dark*, *Dronc*, and *Drice*, or overexpression of the *dominant-negative form of Dronc* (*Dronc^DN^*) failed to suppress the wingless phenotype (Figure 1G, H). These findings support our earlier observation that Dcp-1 can be activated independently of the canonical apoptosome composed of Dronc and Dark to mediate non-lethal functions (Shinoda et al., 2019). Collectively, these results indicate that, at least under overexpression conditions, Dcp-1 activation proceeds independently of the apoptosome, in contrast to that of Drice.

### Dcp-1 and Drice show distinct expression patterns among tissues

Proximal proteins of caspases have emerged as potential regulators of executioner caspase activity (Muramoto et al., 2025). We have previously demonstrated that the proximal protein patterns of Dcp-1 and Drice differ even within the same tissue, such as wing imaginal discs (Shinoda et al., 2019). Based on this, we sought to identify Dcp-1-specific regulatory factors among its proximal proteins. First, we examined the tissue distribution of *Drosophila* caspases using previously established *Caspase::V5::TurboID* knock-in fly lines (Shinoda et al., 2019). Expression patterns were analyzed in larval tissues, including the salivary glands, fat bodies, and brains (Figure 2–figure supplement 1A), with confirmed expression in wing imaginal discs (Figure 2A) as well as in adult tissues, including the thoraxes, ovaries, and testes (Figure 2–figure supplement 1B). Drice was ubiquitously expressed in all examined tissues. In contrast, Dcp-1 expression was restricted to the larval brains, imaginal discs and adult ovaries and testes.

**Figure 2.**
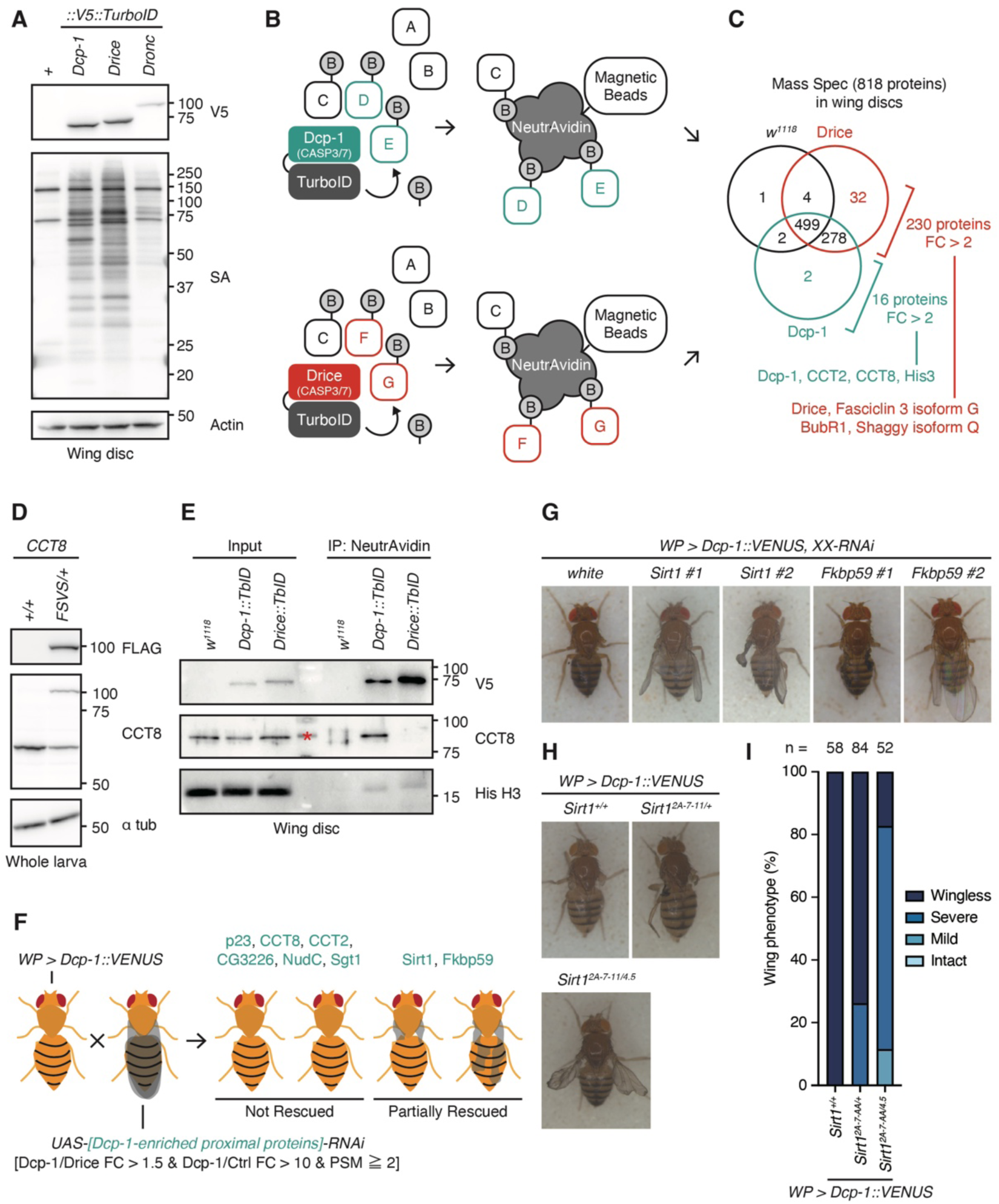
Dcp-1-enriched proximal proteins are required for Dcp-1 activation. (A) Western blotting of expression of C-terminally V5::TurboID knocked-in tagged caspases and biotinylation patterns of their proximal proteins detected by streptavidin (SA) in larval wing imaginal discs. (B) A schematic diagram of the TurboID-mediated identification of proximal proteins. Dcp-1-and Drice-proximal proteins (Protein C, D, E, F, G) are promiscuously labeled *in vivo* with the administration of biotin. Then, biotinylated proteins are purified using NeutrAvidin magnetic beads and are subsequently analyzed by mass spectrometry. (C) A summary of mass spectrometry analysis of proteins in the larval imaginal discs. Among 818 proteins identified, 230 proteins were detected as highly enriched proteins in Drice::V5::TurboID flies (FC [Drice::V5::TurboID/Dcp-1::V5::TurboID] > 2) and 16 proteins were detected as highly enriched proteins in Dcp-1::V5::TurboID flies (FC [Dcp-1::V5::TurboID/Drice::V5::TurboID] > 2). (D) Western blotting validation of the anti-CCT8 antibody in whole larval lysates without the digestive tract and fat body. The anti-CCT8 antibody detects both endogenous CCT8 (MW: 59.4 kDa) and 3xFLAG::StrepII::VENUS::StrepII (FSVS)-tagged CCT8 (MW: 91.1 kDa). (E) Western blotting of endogenous CCT8 expressed in larval wing imaginal disc. Dcp-1-proximal and Drice-proximal proteins biotinylated by C-terminally knocked-in tagged V5::TurboID are purified using NeutrAvidin. The red asterisk represents a non-specific band in the marker lane. (F) Schematic diagram of the RNAi screen targeting eight Dcp-1-enriched proximal proteins. The *WP > Dcp-1::VENUS* line was crossed with each *UAS-RNAi* line, and wing phenotypes of the F1 progeny were scored. (G) Representative images of female wing phenotypes from *Dcp-1::VENUS* overexpression driven by *WP-Gal4* with knockdown of Dcp-1-enriched proximal proteins. “XX” denotes the knocked-down gene, and “#1” and “#2” indicate independent RNAi lines. (H) Representative images of female wing phenotypes upon *Dcp-1::VENUS* overexpression driven by *WP-Gal4* in the *Sirt1* null mutant background. (I) Quantification of (H). Each wing is manually classified into four (wingless, severe, mild, and intact) categories. Sample sizes are shown in the figure.

### Drosophila executioner caspases associate with distinct sets of proximal proteins

Next, we compared the proximal protein profiles of *Drosophila* executioner caspases, based on our previous observation that Dcp-1 and Drice are associated with distinct sets of proximal proteins, potentially reflecting differences in their activation mechanisms (Shinoda et al., 2019). Streptavidin staining, used to visualize proteins biotinylated via TurboID-mediated proximity labeling, revealed distinct patterns of Dcp-1 and Drice expression across tissues (Figure 2A, Figure 2–figure supplement 1A, B). To directly compare their proximal landscapes in the same cellular context, we focused on the imaginal discs. NeutrAvidin-based purification of biotinylated proteins, followed by MS, identified 818 proteins (Figure 2B, C). As expected, the proximal proteomes of Dcp-1 and Drice were distinct. BubR1 and Fasciclin 3 isoform G, previously identified as Drice-proximal in adult brain and ovary tissues (Muramoto et al., 2025; Shinoda et al., 2023), were enriched in the Drice-associated proteome (Figure 2C, Table S1), suggesting that Drice maintains its proximity to specific partners across tissues. Unlike overexpressed Dcp-1, endogenous Dcp-1 expressed in wing imaginal discs exists in its full-length form (Figure 2–figure supplement 2A), suggesting that the identified Dcp-1 proximal proteins are associated with the full-length pro-form. Consistently, DIAP-1, which is shown to interact with Dcp-1 and Drice only after exposure of the IAP-binding motif (IBM) at the neo N-terminus of the large executioner caspase subunit following cleavage (Tenev et al., 2005), is not identified in the Dcp-1 proximal protein list (Figure 2C, Table S1). To validate the TurboID-MS result, we compared our Dcp-1 proximal protein list with the previously reported 24 Dcp-1 interactors identified in *Drosophila* l(2)mbn cultured cells by immunoaffinity purification (IAP) followed by MS (Choutka et al., 2017). Although the cell types and methods are different, 15 proteins are identified in both analyses (Figure 2–figure supplement 2B, Table S2). Among these, 8 proteins were identified as highly enriched Dcp-1 proximal proteins (Dcp-1/Control fold change ≧ 2; Figure 2–figure supplement 2B, Table S2). Importantly, SesB, one of the best-characterized Dcp-1 interactors located in mitochondria (DeVorkin et al., 2014), was also identified in our TurboID-MS dataset (Figure 2–figure supplement 2B, Table S2), indicating that our TurboID-MS analysis successfully identified Dcp-1 proximal proteins. Among the top Dcp-1-enriched proximal proteins was the chaperonin containing TCP1 subunit 8 (CCT8). To validate these findings, we generated an antibody against CCT8 (Figure 2D). Western blotting confirmed the presence of CCT8 in Dcp-1::V5::TurboID samples but not in Drice::V5::TurboID samples (Figure 2E). Taken together, these results show that, even when expressed at comparable levels in the same tissue, Dcp-1 and Drice are associated with distinct sets of proximal proteins, which is consistent with their differential regulation and functional specificity.

### Screening of Dcp-1 proximal proteins identifies Sirt1 as a Dcp-1 activator

To identify potential regulators of Dcp-1 activation, we performed an RNAi screen targeting eight genes that were selectively proximal to Dcp-1 relative to Drice and wild-type controls (Dcp-1/Drice fold change > 1.5, Dcp-1/Control fold change > 10 and PSM ≧ 2; Table S1), using two independent RNAi lines per gene (Figure 2F). Under normal conditions, RNAi-mediated knockdown of the chaperonin components *CCT2* and *CCT8* resulted in a wingless phenotype, whereas knockdown of other genes had no effect (Figure 2–figure supplement 3A). Under *Dcp-*1 overexpression conditions, knockdown of *Fkbp59* or *Sirt1* partially suppressed Dcp-1-induced cell death (Figure 2G, Figure 2–figure supplement 3B). The requirement for Sirt1 was further confirmed using a trans-heteroallelic combination of two previously described *Sirt1* deletion alleles, *Sirt1^2A-7-11^*(Furuyama et al., 2004; Xie and Golic, 2004) and *Sirt1^4.5^* (Newman et al., 2002), which significantly rescued the wingless phenotype (Figure 2H, I). These results indicate that the Dcp-1-proximal protein Sirt1 is required for Dcp-1 activation upon overexpression.

### Dcp-1 activation specifically depends on Debcl, Buffy, and autophagy

Sirt1, also known as dSir2, is a lysine deacetylase that promotes the expression of autophagy-related genes via Atg8a deacetylation under starvation (Jacomin et al., 2020). Multiple studies have linked Dcp-1 to the regulation of autophagy. Dcp-1, together with the Bcl-2 family proteins Debcl and Buffy, but not Drice, has been shown to be required for starvation-induced autophagy in *Drosophila* l(2)mbn cells (Hou et al., 2008), and Dcp-1 is also required for autophagy in the ovary (Choutka et al., 2017; DeVorkin et al., 2014; Hou et al., 2008). More recently, the BH3-only protein Sayonara (Synr) was reported to induce Dcp-1-dependent cell death, which is regulated by Debcl, Buffy, and autophagy (Ikegawa et al., 2023). In contrast, a gain-of-function modifier screen revealed that autophagy antagonizes Dcp-1-induced cell death (Kim et al., 2010) and that Dcp-1 suppresses autophagy through cleavage-mediated inactivation of Acinus (Haberman et al., 2010; Nandi et al., 2014).

Based on these reports, we examined whether Synr, Debcl, Buffy, and core autophagy components regulate Dcp-1 activation. Knockdown of *Synr* did not suppress Dcp-1-induced cell death, whereas knockdown of *Debcl* or *Buffy* significantly rescued this wingless phenotype (Figure 3A, B). Similarly, knockdown of the autophagy-related genes *Atg2* and *Atg8a* suppressed Dcp-1-induced cell death (Figure 3A, B), indicating that Debcl, Buffy, and autophagy are required for Dcp-1 activation. Of note, knockdown of any of the genes tested did not affect wing morphology in the absence of *Dcp-1* overexpression (Figure 3–figure supplement 1A), suggesting the specific requirement for Dcp-1 activation. We further performed knockdown of autophagy-related genes including *FIP200/Atg17*, which is required for the initiation of autophagosome formation; *Atg9*, which is required for autophagosomal membrane nucleation; *Atg5*, which is required for autophagosomal membrane expansion through the Atg12-Atg5-Atg16 ubiquitin-like conjugation system; and *Stx17*, which is required for autophagosome-lysosome fusion (Umargamwala et al., 2024). While knockdown of these genes alone did not affect wing morphology in the absence of *Dcp-1* overexpression (Figure 3–figure supplement 1B), knockdown of each of these genes suppressed the Dcp-1-induced wing ablation phenotype (Figure 3–figure supplement 1C), supporting a requirement for canonical autophagy components across multiple stages of autophagosome biogenesis in Dcp-1 activation. Consistent with the requirement of autophagy for Dcp-1 activation, induction of autophagy, as judged by the accumulation of the formation of autophagosomes marked by mCherry-tagged Atg8a puncta, was observed upon *Dcp-1* overexpression (Figure 3C). These results further support the idea that autophagy promotes Dcp-1 activation.

**Figure 3.**
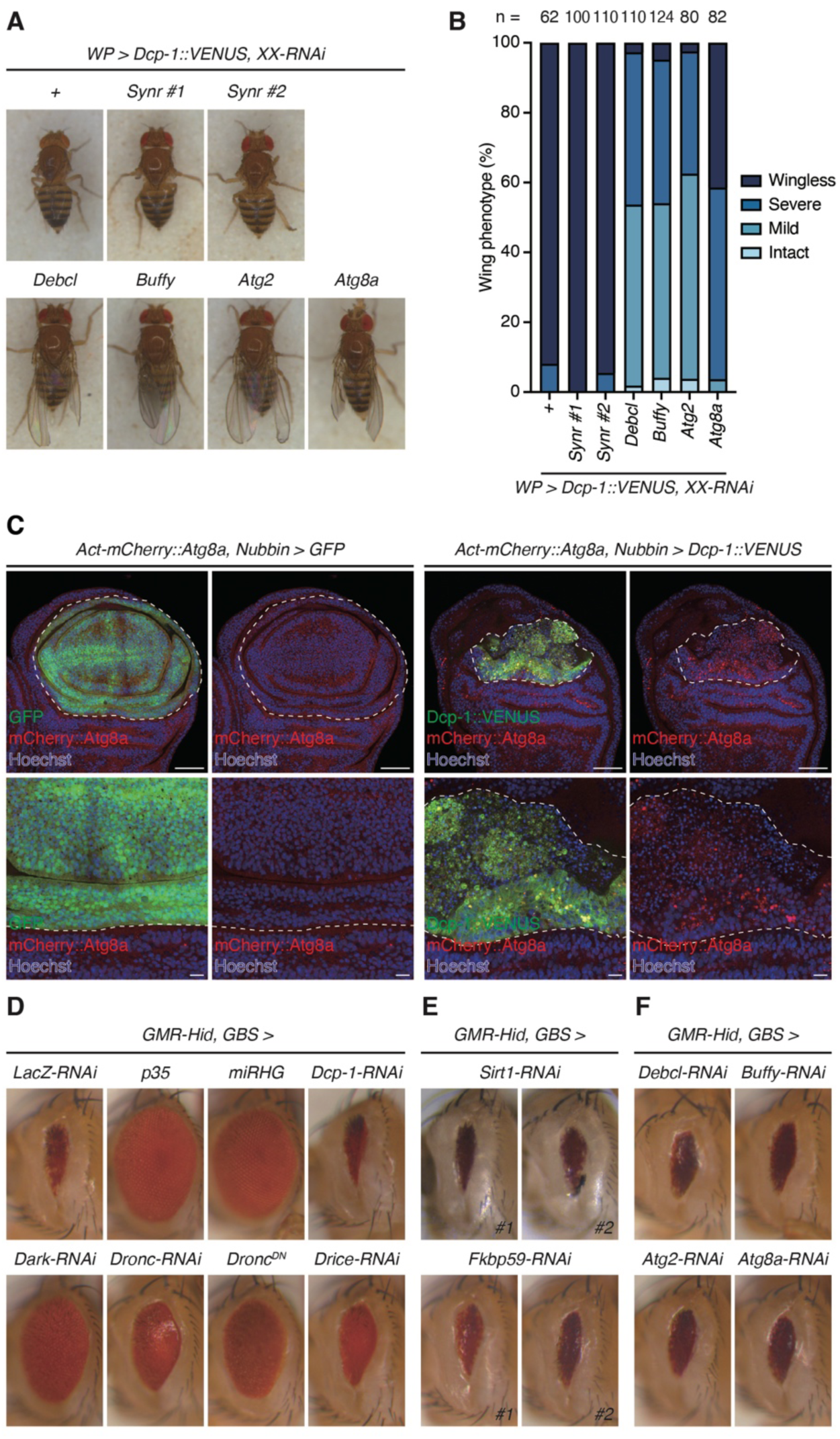
Debcl, Buffy, and autophagy are specifically required for *Dcp-1* overexpression-induced apoptosis. (A) Representative images of female wing phenotypes from *Dcp-1::VENUS* overexpression driven by *WP-Gal4* with the genetic perturbation of *Synr*, *Debcl*, *Buffy,* and autophagy. “XX” denotes the knocked-down gene, and “#1” and “#2” indicate independent RNAi lines. (B) Quantification of (A). “XX” denotes the knocked-down gene, and “#1” and “#2” indicate independent RNAi lines. Each wing is manually classified into four (wingless, severe, mild, and intact) categories. Sample sizes are shown in the figure. (C) Representative images of mCherry::Atg8a puncta (autophagosome) accumulation (red) in larval wing imaginal discs upon *VENUS-tagged Dcp-1* (green) overexpression using *WP-Gal4*. Nuclei are visualized by Hoechst 33342 (blue). Scale bar: 50 µm (top) and 10 µm (bottom). The contrast of GFP and Dcp-1::VENUS signals are adjusted differently to enhance visibility. (D) Representative images of female eye phenotypes upon *Hid* overexpression with the genetic perturbation of canonical apoptosis signaling. (E) Representative images of female eye phenotypes upon *Hid* overexpression with the genetic perturbation of *Sirt1* or *Fkbp59*. “#1” and “#2” indicate independent RNAi lines. (F) Representative images of female eye phenotypes upon *Hid* overexpression with the genetic perturbation of *Debcl*, *Buffy*, and autophagy.

To assess the specificity of these regulators, we analyzed the eye ablation phenotype caused by overexpression of *Hid*, which triggers apoptosis via canonical apoptotic signaling. As expected, this phenotype was rescued by *p35* expression or knockdown of *RHG* genes and partially suppressed by inhibition of the Dark-, Dronc-, and Drice-dependent apoptosome pathways (Figure 3D), confirming its dependence on canonical apoptosis (Figure 1E). In contrast, *Hid*-induced apoptosis was not rescued by RNAi against *Dcp-1* (Figure 3D), nor by knockdown of *Sirt1* or *Fkbp59* (Figure 3E). Furthermore, knockdown of *Debcl*, *Buffy*, and autophagy genes failed to rescue *Hid*-induced apoptosis (Figure 3F). Taken together, these results indicate that Sirt1, Debcl, Buffy, and autophagy components are not general regulators of apoptosis but function as specific activators of Dcp-1.

### Cleaved Dcp-1 specifically interacts with a giant-IAP Bruce

Next, we examined the molecular link between autophagy and Dcp-1 activation. Caspase activation is negatively regulated by IAPs. In *Drosophila*, DIAP-1 suppresses caspase activity through ubiquitination (Ditzel et al., 2008) (Figure 1E). In addition to DIAP-1, *Drosophila* has another giant IAP, Bruce, which is degraded via autophagy during oogenesis (Nezis et al., 2010). Bruce inhibits autophagy under nutrient-rich conditions (Hou et al., 2008), suggesting mutual negative regulation of Bruce and autophagy. Bruce selectively inhibits reaper-induced cell death, but not hid-induced apoptosis, suggesting a degree of pathway specificity (Domingues and Ryoo, 2012). These observations led us to focus on Bruce as a potential selective regulator of Dcp-1 activation.

The structure of human Bruce/BIRC6 has recently been resolved using cryo-electron microscopy (Dietz et al., 2023; Ehrmann et al., 2023; Hunkeler et al., 2023; Liu et al., 2024), revealing several conserved domains: linker domain (LD), baculoviral IAP repeat (BIR), WD40, β-X1, β-X2, carbohydrate-binding module (CBM), ubiquitin-like (UBL), and ubiquitin-conjugating (UBC) (Figure 4A). Human Bruce/BIRC6 ubiquitinates the activated executioner caspases caspase-3 and caspase-7, but not their non-processed proforms *in vitro* (Dietz et al., 2023). The BIR domain mediates interactions with ubiquitination substrates by recognizing N-terminal IAP-binding motifs (Berthelet and Dubrez, 2013). The consensus IBM sequence is defined as A(K/T/V/I)(P/A/E)(F/E/I/S/Y) (Vaux and Silke, 2005). We identified IBM-like sequences at the N-termini of the P20 domains of both Dcp-1 and Drice (Figure 4B, C). To assess the potential interaction between Bruce and these caspases, we used AlphaFold3 to predict the structure of the complexes between Bruce and the IBM-like motifs of Dcp-1 and Drice (Abramson et al., 2024). Consistent with the known architecture of BIRC6, AlphaFold3 predicted that the BIR domain of Bruce interacts with LD, WD40, β-X1, and β-X2 domains (Figure 4–figure supplement 1A, A’, B, B’). For structural prediction, we used a Bruce fragment spanning residues 1–1960, including the BIR domain and its associated Zn²⁺-binding site (Figure 4A), and 15-residue peptides from the Dcp-1 and Drice cleavage sites encompassing their IBM-like motifs (Figure 4B, C). The predicted structures revealed a high-confidence interaction between Dcp-1 and the Bruce BIR domain. All five AlphaFold3 models showed confident pLDDT scores (70–90) for the N-terminal residues of Dcp-1, corresponding to the IBM-like motif (Figure 4D, E). The predicted binding mode resembled the canonical BIR-IBM interaction observed in XIAP BIR3-Smac (Wu et al., 2000) (Figure 4–figure supplement 1C, C’). In contrast, although Drice appeared to bind to Bruce in the predicted models, the pLDDT scores were consistently low (<70) across the interaction interface, including the four N-terminal residues, indicating low confidence in the predicted interaction (Figure 4F, G).

**Figure 4.**
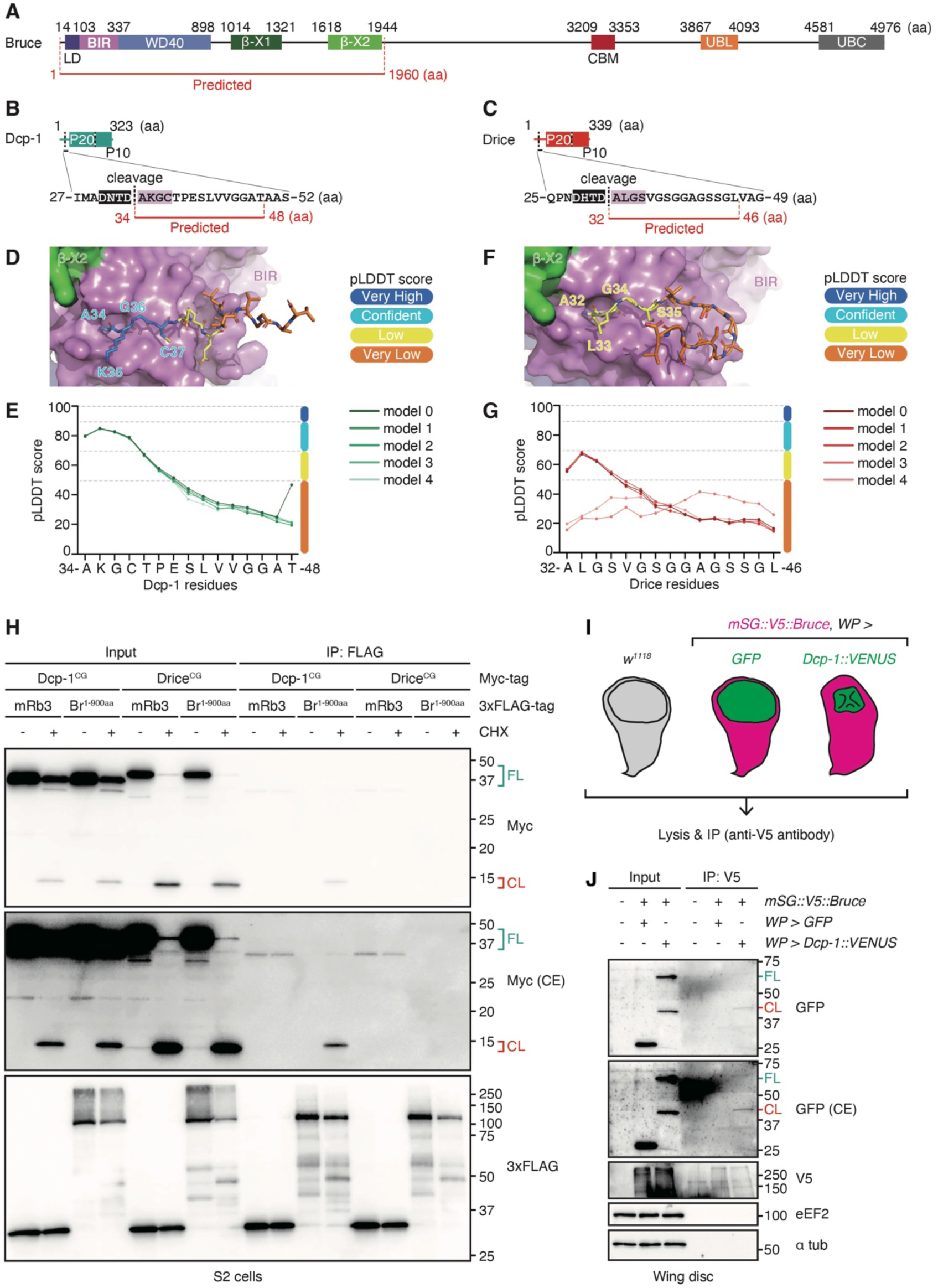
Cleaved Dcp-1, but not Drice, interacts with Bruce. (A) Domain structure of Bruce. LD denotes the linker domain, BIR the Baculoviral IAP repeat, WD40 a WD40 domain, β-X a β-sandwich-like fold, CBM a domain resembling the carbohydrate-binding module family 32, UBL the ubiquitin-like domain, and UBC the E2-E3 hybrid ubiquitin-conjugating domain. The region used for structural prediction using AlphaFold3 is indicated in red. (B) IBM-like sequence in Dcp-1. The propeptide cleavage site of Dcp-1 is indicated by a black dashed line; the caspase recognition motif is highlighted in black; the IBM-like sequence is highlighted in pink; and the region used for AlphaFold3 prediction is shown in red. (C) IBM-like sequence in Drice. The propeptide cleavage site of Drice is indicated by a black dashed line; the caspase recognition motif is highlighted in black; the IBM-like sequence is highlighted in pink; and the region used for AlphaFold3 prediction is shown in red. (D) Predicted complex structure of Bruce BIR and Dcp-1 IBM-like sequence. The AlphaFold3 prediction [model 0] is shown. The Bruce β-X2 and BIR domains are displayed as surfaces in green and pink, respectively. Dcp-1 is shown in stick representation and color-coded according to pLDDT scores, as indicated in the figure. See also panel (E). (E) pLDDT scores of Dcp-1 in the five predicted complex structures with Bruce. For all five AlphaFold3 models, per-residue pLDDT scores of Dcp-1 are plotted. Prediction confidence is color-coded as follows: very high (pLDDT > 90) in dark blue, confident (90 > pLDDT > 70) in light blue, low (70 > pLDDT > 50) in yellow, and very low (pLDDT < 50) in orange. (F) Predicted complex structure of Bruce BIR and Drice IBM-like sequence. The AlphaFold3 prediction [model 0] is shown. The Bruce β-X2 and BIR domains are displayed as surfaces in green and pink, respectively. Drice is shown in stick representation and color-coded according to pLDDT scores, as indicated in the figure. See also panel (G). (G) pLDDT scores of Drice in the five predicted complex structures with Bruce. For all five AlphaFold3 models, per-residue pLDDT scores of Drice are plotted. Prediction confidence is color-coded as follows: very high (pLDDT > 90) in dark blue, confident (90 > pLDDT > 70) in light blue, low (70 > pLDDT > 50) in yellow, and very low (pLDDT < 50) in orange. (H) Co-immunoprecipitation of N-terminally 3xFLAG-tagged N-terminal region of Bruce (Br, 1–900 aa; containing LD, BIR, and WD40 domains) with C-terminally myc-tagged, catalytically inactive Dcp-1 or Drice (catalytic cysteine mutated to glycine; CG) expressed in *Drosophila* S2 cells. N-terminally 3xFLAG-tagged mRuby3 (mRb3) was used as a control. Apoptosis was induced by cycloheximide (CHX) treatment. 3xFLAG-tagged proteins were immunoprecipitated using an anti-3xFLAG antibody. Both full-length (FL) and cleaved (CL) caspases are detected using anti-Myc antibody. For the anti-Myc blot, a contrast-enhanced (CE) image is shown below. (I) Schematic diagram of wing imaginal disc samples used for co-immunoprecipitation to analyze the interaction between endogenously expressed N-terminally mStayGold::V5-tag knocked in-tagged Bruce (magenta) with C-terminally VENUS-tagged Dcp-1 (green). (J) Co-immunoprecipitation of endogenously expressed N-terminally mStayGold::V5-tag knocked in-tagged Bruce with C-terminally VENUS-tagged Dcp-1 overexpressed in wing imaginal disc using *WP-Gal4*. GFP overexpressed in wing imaginal disc using *WP-Gal4* was used as a control. V5-tagged proteins were immunoprecipitated using an anti-V5-tag antibody. Both full-length (FL) Dcp-1::VENUS (MW: 62.7 kDa) and cleaved (CL) Dcp-1::VENUS (MW: 39.1 kDa) are detected using anti-GFP antibody. For the anti-GFP blot, a contrast-enhanced (CE) image is shown below.

To validate these predictions, we performed co-immunoprecipitation assays on *Drosophila* S2 cells. We found that only the cleaved form of Dcp-1, which forms a head-to-tail dimer composed of the P20 and P10 domains (Julien and Wells, 2017), and not the full-length protein, interacted with the N-terminal BIR domain of Bruce following apoptosis induction (Figure 4H), consistent with the observation that human Bruce/BIRC6 only targets cleaved executioner caspases for ubiquitination *in vitro* (Dietz et al., 2023). Conversely, neither the cleaved nor full-length forms of Drice showed detectable interactions with the Bruce BIR domain (Figure 4H), corroborating the AlphaFold3 predictions.

To further examine whether endogenous Bruce interact with activated Dcp-1, we generated CRISPR/Cas9-mediated knock-in flies in which an mStayGold::C4 linker::V5-tag (Ando et al., 2024) was inserted into the N-terminus of Bruce (Figure 4–figure supplement 2A). Bruce has been reported to function in oogenesis and spermatogenesis in the ovaries and testes, respectively (Arama et al., 2007, 2003; Hou et al., 2008; Kaplan et al., 2010; Nezis et al., 2010). However, its expression in other tissues has not been thoroughly characterized. Using the knocked-in line, we confirmed that Bruce was expressed in the ovaries (Figure 4–figure supplement 2B) and testes (Figure 4–figure supplement 2C), consistent with previous reports (Arama et al., 2007, 2003; Hou et al., 2008; Kaplan et al., 2010; Nezis et al., 2010). Western blot analysis revealed Bruce expression in both adult males and females (Figure 4–figure supplement 2D). We also detected Bruce expression in wing imaginal discs (Figure 4–figure supplement 2E). This signal was abolished by the RNAi-mediated knockdown of *Bruce*, confirming the specificity of the signal (Figure 4–figure supplement 2F). These findings indicate that Bruce not only functions in the gonads but can also exert Dcp-1 regulatory functions in wing imaginal discs. We overexpressed *Dcp-1::VENUS* using *WP-Gal4* in the background of the established *mStayGold::C4 linker::V5-tag* knock-in-tagged *Bruce* and conducted co-immunoprecipitation analysis (Figure 4I). Following immunoprecipitation with an anti-V5 tag antibody, we found that cleaved Dcp-1 signal was enriched by co-immunoprecipitation (Figure 4J), suggesting that full-length Bruce interacts with activated Dcp-1. Consistently, Bruce was not identified as a proximal protein of full-length Dcp-1 in our TurboID-MS analysis (Figure 2C, Table S1). Taken together, these findings suggest that Bruce specifically suppresses activated Dcp-1 but not Drice and serves as a branching point for a Dcp-1-specific, non-canonical regulatory pathway.

### Bruce specifically suppresses Dcp-1 overexpression-induced cell death

Next, we investigated whether Bruce regulated Dcp-1 activity. Upon overexpression of *Bruce* (Figure 5A), we observed marked suppression of Dcp-1-induced cell death (Figure 5B, C). Although *DIAP-1* overexpression also inhibited Dcp-1 activation, this suppressive effect was stronger following *Bruce* overexpression (Figure 5B, C). We examined the specificity of Bruce in the regulation of other forms of cell death. As mentioned earlier, the BH3-only protein Synr induces Debcl/Buffy, autophagy and Dcp-1-dependent cell death (Ikegawa et al., 2023). To investigate whether Bruce was involved in Debcl/Buffy-autophagy-Dcp-1-mediated cell death, we examined the role of Bruce in Synr-induced cell death. Overexpression of *Bruce*, but not *DIAP-1*, specifically suppressed wrinkled wing phenotype caused by Synr-induced cell death (Figure 5D). In contrast, *Bruce* overexpression failed to rescue the eye ablation phenotype independent of autophagy caused by *Hid* overexpression, whereas DIAP-1 fully rescued this phenotype (Figure 5E). Taken together, these results suggest that Bruce and DIAP-1 regulate distinct cell death pathways, with Bruce acting as a selective inhibitor of autophagy-regulated Dcp-1- and Synr-dependent cell death.

**Figure 5.**
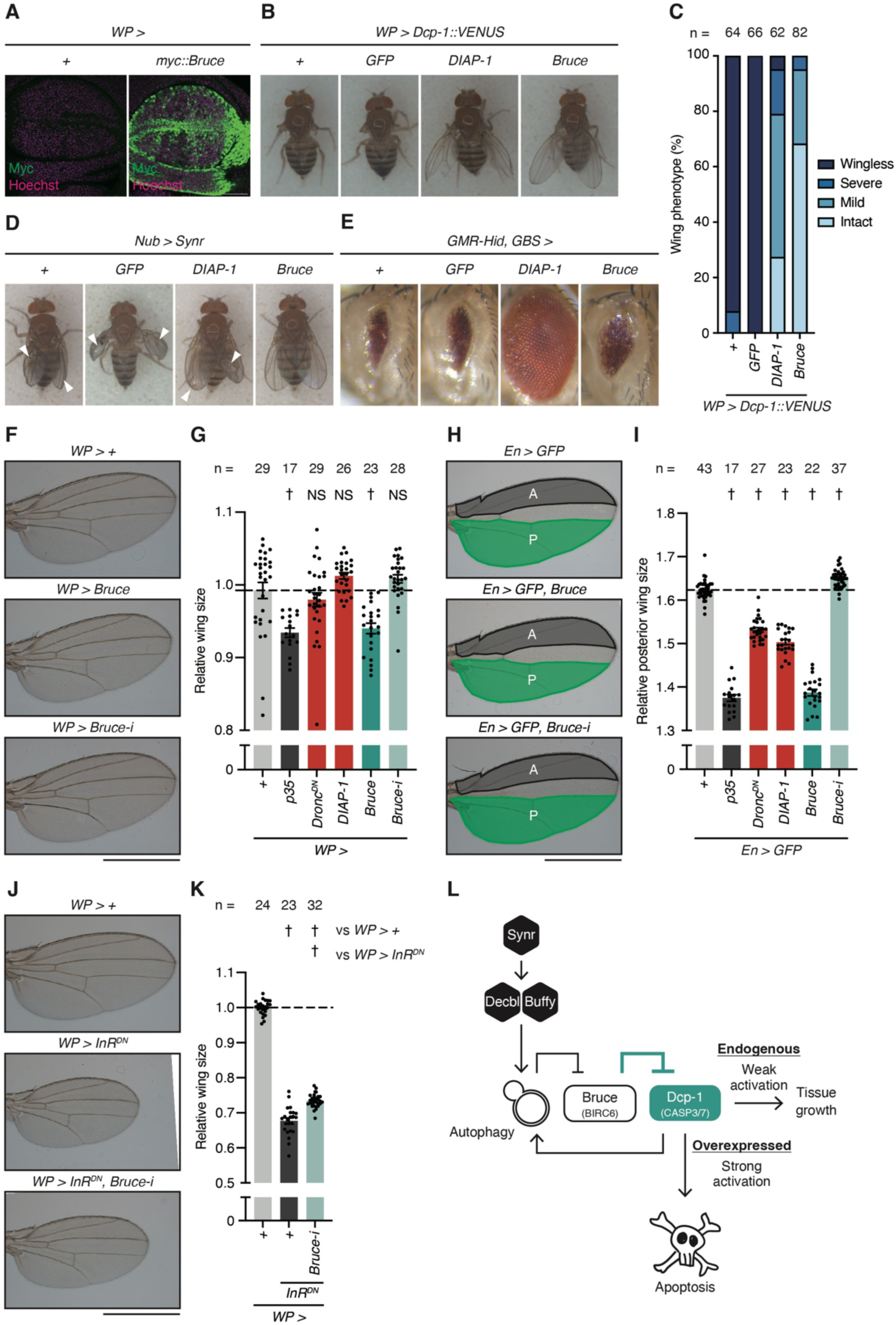
Bruce selectively suppresses Dcp-1 activity and wing tissue growth. (A) Expression pattern of overexpressed *myc::Bruce* (green) by using *WP-Gal4* in the wing imaginal discs. Nuclei are visualized using Hoechst 33342 (magenta). Scale bar: 50 µm. (B) Representative images of female wing phenotypes from *Dcp-1::VENUS* overexpression driven by *WP-Gal4* with *DIAP-1* or *Bruce* overexpression. (C) Quantification of (B). Each wing is manually classified into four (wingless, severe, mild, and intact) categories. Sample sizes are shown in the figure. (D) Representative images of female wing phenotypes upon *Synr* overexpression driven by *Nubbin-Gal4* with *DIAP-1* or *Bruce* overexpression. Arrowheads indicate wrinkled wings. (E) Representative images of female eye phenotypes upon *Hid* overexpression with *DIAP-1* or *Bruce* overexpression. (F) Representative images of female wing upon *Bruce* or *Bruce-RNAi* (*Bruce-i*) overexpression using *WP-Gal4*. Scale bar: 1 mm. (G) Quantification of relative wing size normalized to the corresponding *No-Gal4* control. Data are presented as mean ± SEM. *p*-values were calculated using one-way analysis of variance (ANOVA) followed by Dunnett’s multiple-comparison test against no UAS control. NS, *p* ≧ 0.05; †, *p* < 0.05. Sample sizes are shown in the figure. (H) Representative images of female wing upon *Bruce* or *Bruce-RNAi* (*Bruce-i*) overexpression using *Engrailed-Gal4* (*En-Gal4*). The grey area (labelled as A) represents the anterior part, and the green area (labelled as P) represents the posterior part of the wing. Scale bar: 1 mm. (I) Quantification of relative posterior part of wing size normalized to corresponding anterior part of wing size. Data are presented as mean ± SEM. *p*-values were calculated using one-way ANOVA followed by Dunnett’s multiple comparison test against no UAS control. †, *p* < 0.05. Sample sizes are shown in the figure. (J) Representative images of female wing upon *InR^DN^* and *Bruce-RNAi* (*Bruce-i*) overexpression using *WP-Gal4*. Scale bar: 1 mm. (K) Quantification of relative wing size normalized to control (*WP > +*). Data are presented as mean ± SEM. p-values were calculated using one-way ANOVA followed by Sidak’s multiple comparison test for selected pairs. †, *p* < 0.05. Sample sizes are shown in the figure. (L) Schematic diagram of autophagy-facilitated alternative caspase activation pathway in *Drosophila* mediated by autophagy-Bruce axis.

### Bruce suppresses wing tissue growth

We have previously reported that Dcp-1 activity can promote imaginal tissue growth independent of the core apoptosis signaling pathway, including Dronc (Shinoda et al., 2019). To investigate the role of Bruce in imaginal tissue growth, we analyzed its effect on wing imaginal discs using *WP-Gal4*, which drives expression throughout the wing pouch. Overexpression of *p35* suppressed wing tissue growth (Figure 5F, G), whereas inhibition of canonical apoptotic signaling by *Dronc^DN^* and *DIAP-1* had no effect (Figure 5F, G). In contrast, *Bruce* overexpression suppressed wing growth to an extent similar to *p35* overexpression, whereas *Bruce* knockdown tended to increase wing size, although the difference was not statistically significant (Figure 5F, G). We confirmed these results using *Engrailed-Gal4*, which is expressed in the posterior compartment of the wing imaginal disc. *p35* overexpression suppressed wing tissue growth (Figure 5H, I). In this context, the inhibition of canonical apoptosis signaling by *Dronc^DN^* and *DIAP-1* had a minor but negative effect on wing size (Figure 5H, I). *Bruce* overexpression suppressed wing growth, whereas *Bruce* knockdown increased wing size (Figure 5H, I), indicating that Bruce negatively regulates wing tissue growth. We also found that knockdown of *Debcl* and *Buffy* resulted in reduced wing size (Figure 5–figure supplement 1A), suggesting that Debcl and Buffy also participate in this process. Finally, to enhance the sensitivity of the assay, we overexpressed the dominant-negative form of the insulin receptor (*InR^DN^*) using *WP-Gal4*, which significantly reduced wing size (Figure 5J, K). Under this sensitized condition, simultaneous *Bruce* knockdown significantly enhanced wing tissue growth (Figure 5J, K). Overall, these results indicate that Bruce suppresses wing tissue growth.

## Discussion

### Autophagy-facilitated alternative caspase activation pathway

In this study, we identified a distinct regulatory mechanism governing the activation of the *Drosophila* executioner caspase, Dcp-1, in sharp contrast to its paralog Drice. Although both caspases share structural similarities and are expressed in overlapping tissues, Dcp-1, but not Drice, is activated upon overexpression in an apoptosome-independent, autophagy-regulated manner. Previous studies have implicated Dcp-1 in the induction of autophagy (Choutka et al., 2017; DeVorkin et al., 2014; Hou et al., 2008). Our findings extend this view by showing that autophagy is required for Dcp-1 activation, suggesting a positive feedback loop between Dcp-1 and autophagy during apoptosis. Although the precise downstream targets of the Bcl-2 family proteins, Debcl and Buffy, remain unknown, their roles in autophagy regulation (Hou et al., 2008) suggest that they influence Dcp-1 activity through autophagic pathways. Additionally, we identified Bruce, a giant inhibitor of apoptotic proteins, as a Dcp-1-specific suppressor. Functionally, Bruce limits wing tissue growth, suggesting that Dcp-1 activation contributes to growth regulation in a non-lethal manner (Shinoda et al., 2019). Taken together, our results demonstrate that the regulatory mechanisms of Dcp-1 and Drice diverge at the level of IAPs and that Dcp-1 activation is regulated by autophagy, enabling specific non-lethal functions that promote tissue growth independent of canonical apoptotic signaling (Figure 5L). It is important to note that Dcp-1 can also suppress autophagy through cleavage of Acinus (Haberman et al., 2010; Nandi et al., 2014). Although this may seem contradictory, the ability of caspases to function in a non-lethal manner requires strict prevention of excessive activation. Such suppression may represent a key mechanism that enables non-lethal caspase activity.

We have previously shown that another executioner caspase, Decay, also participates in promoting imaginal tissue growth (Shinoda et al., 2019). However, the upstream regulatory mechanisms of Decay remain unknown. Importantly, Decay has been shown to mediate Hid-induced cell death in the DIAP1- and apoptosome-independent manner in differentiating photoreceptors and accessory cells of the eye (Leulier et al., 2006). In addition, although not required for cell death, Decay accounts for most of the caspase activity during metamorphic midgut programmed cell death, which is executed by autophagy (Denton et al., 2009). Thus, similar to Dcp-1, Decay might be an executioner caspase that can be regulated independently of the canonical apoptosome-mediated pathway, potentially involving autophagy-Bruce axis, and thereby contributing to the regulation of tissue growth.

It is also important to understand how cells determine the usage of canonical versus alternative caspase activation pathways depending on physiological or environmental conditions. In mammals, two major apoptotic pathways are activated by distinct initiator caspases, caspase-8 and caspase-9, in response to different upstream signals (Newton et al., 2024). Similarly, in *Drosophila*, while most developmental cell death is driven by canonical apoptotic signaling involving Dronc (Xu et al., 2005), alternative caspase activation pathways may be preferentially engaged in response to environmental stimuli, such as starvation or heat shock.

### Initiator caspase-independent activation mechanism for Dcp-1

The mechanism by which Dcp-1 is activated by itself remains an open and important question. In contrast to Drice, Dcp-1 is activated upon overexpression in an apoptosome-independent but autophagy-dependent manner. Unlike initiator caspases, executioner caspases such as Dcp-1 generally lack long N-terminal prodomains and require proteolytic cleavage to become catalytically active. Proximity-labeling analysis revealed that Dcp-1 was predominantly associated with the chaperonin subunit CCT8 (Figure 2E). Given that the overexpression of *Dcp-1* increases its local concentration for activation, we propose that its accumulation in the chaperonin complex may serve as a local scaffold for caspase activation. This model implies a spatially restricted mode of regulation in which caspase activity is compartmentalized within the cell. In this model, we speculated that Dcp-1 activation occurs sporadically in subcellular regions enriched in activators or scaffolding molecules such as CCT8. Under these conditions, Dcp-1 activation may need to be tightly regulated to prevent its unintended proteolytic activity. Supporting this notion, we proposed that Bruce functions as a basal suppressor of Dcp-1 in non-lethal contexts, restraining caspase activity when full apoptotic signaling is not engaged.

### Distinct roles of evolutionarily multiplicated executioner caspases

Caspases are essential cysteine proteases that execute apoptotic cell death and are conserved from *C. elegans* to mammals (Shinoda and Miura, 2024). Multiple executioner caspases exist, including caspase-3 and caspase-7 in mammals and Dcp-1 and Drice in *Drosophila*. Although these enzymes are often functionally redundant in apoptosis, growing evidence indicates that each executioner caspase harbors distinct substrate specificities and fulfills non-overlapping roles (Boucher et al., 2012; Desroches and Denault, 2019; Shinoda et al., 2023, 2019; Walsh et al., 2008). In mammals, caspase-3 and caspase-7 exhibit high sequence similarity but differ in their substrate preferences. Caspase-7, for instance, contains polybasic residues in its exosite that enhance its ability to cleave RNA-binding proteins, a function not observed in caspase-3 (Boucher et al., 2012). Mechanistically, this exosite enables caspase-7 to interact directly with RNA, thereby promoting selective proteolysis of RNA-associated substrates (Desroches and Denault, 2019). Furthermore, we have recently demonstrated that caspase-7 is essential for early-peaking caspase activity at the plasma membrane during apoptosis, a process mediated by its N-terminal intrinsically disordered region, which facilitates rapid efferocytosis of apoptotic cells (Taira et al., 2025). In this study, we demonstrated that *Drosophila* executioner caspases Dcp-1 and Drice are also subject to distinct activation mechanisms for the specific non-lethal function of tissue growth. These findings underscore that the evolutionary expansion of executioner caspases does not merely provide functional redundancy for apoptotic execution but rather confers unique biochemical properties and regulatory roles under distinct cellular contexts. This diversification allows executioner caspases to participate in both lethal and non-lethal cellular processes, highlighting their broader functional repertoire beyond canonical apoptosis.

## Methods

### Fly strains and rearing conditions

We raised the flies in standard *Drosophila* medium at 25°C. For in vivo biotin labeling experiments, flies were raised in standard *Drosophila* medium supplemented with 100 μM (+)-biotin (#029-08713, WAKO). The fly strains used in this study are listed in Table S3. Genotypes corresponding to each figure are shown in Table S4.

### Intrinsically disordered region prediction

Amino acid sequences were obtained from the FlyBase database [https://flybase.org/]. Disordered regions were predicted using the PSIPRED protein sequence analysis workbench in DISOPRED3 [http://bioinf.cs.ucl.ac.uk/psipred/].

### Adult wings and eyes phenotypes scoring

Adult female flies were anesthetized using CO₂. The wing and eye images were acquired using a Leica MZ16F fluorescent stereomicroscope (Leica Microsystems) equipped with a Leica DFC450 C CCD camera. Each wing was manually classified into one of four categories: *Intact* (no defect), *Mild* (≧50% of wing area intact), *Severe* (<50% but ≧5% of wing area intact), and *Wingless* (<5% of wing area intact).

### Molecular cloning

For *pAc5-Caspases-myc-mNeonGreen*, the coding sequences of *Dcp-1* and *Drice* were amplified using PCR from the corresponding *pUASz-Caspase::V5::TurboID* plasmids (Muramoto et al., 2025). The coding sequence of myc-tagged mNeonGreen was PCR-amplified from *pUASz-Drice::myc::mNeonGreen plasmid* (Muramoto et al., 2025). The two fragments were ligated into EcoRI/XhoI-digested *pAc5_STABLE2_neo* (González et al., 2011) using In-Fusion (TaKaRa, Z9648N).

For *pCold I-EcoRI-CCT8-EcoRI*, the coding sequence of *CCT8* was PCR-amplified from *Drosophila* cDNA, and the fragment was ligated into the *Eco*RI-digested *pCold I DNA* vector (#3361, TaKaRa) using In-Fusion.

For *pAc5-Caspases^CG^-myc*, the coding sequences of *Dcp-1* and *Drice* were PCR amplified from the corresponding *pAc5-Caspases-myc-mNeonGreen* plasmids. The fragments were ligated into EcoRI/BamHI-digested *pAc5_STABLE2_neo* (González et al., 2011) using In-Fusion. For pAct-Bruce^1-900a^a-3xFLAG, the first 900 aa of the coding sequences of *Bruce* were PCR-amplified from *Drosophila* cDNA, and the fragment was ligated into the *Eco*RI-digested *pAc5-Fasciclin 3 isoform A-3xFLAG* plasmids (Muramoto et al., 2025) using In-Fusion. For *pAc5-3xFLAG-mRuby*3, *Drosophila* codon-optimized mRuby3 was synthesized (Integrated DNA Technologies, Coralville, IA, USA) and ligated into the BamHI-digested *pAc5-3xFLAG-Bub3* plasmids (Shinoda et al., 2023) using In-Fusion.

For mStayGold::C4 linker::V5-tag knock-in for Bruce, the guide RNA was identified using the CRISPR Optimal Target Finder tool available on flyCRISPR [http://flycrispr.molbio.wisc.edu/]. The DNA fragments for the guide RNA were subcloned into BbsI-digested pDCC6 vectors (#59985, Addgene) (Gokcezade et al., 2014) to establish the *pDCC6-Bruce-sgRNA* plasmid. The following primers were annealed to generate the DNA fragments for guide RNAs: 5′-CTTCGCTTGCTTGAGGACGCGCTGC-3′ and 5′-AAACGCAGCGCGTCCTCAAGCAAGC-3′. For the generation of the homology-directed repair template (*pBac-mStayGold-c4-V5-Bruce-donor-3xP3-dsRed-polyA*), DNA fragments of the 5′ homology arm were assembled into the BsaI site, and the DNA fragment of the 3′ homology arm with mStayGold::C4 linker::V5-tag was assembled into the SapI site of pBac[3xP3-DsRed_polyA_Scarless_TK] (Shinoda et al., 2019) using In-Fusion. The PAM sequence (TGG) of the gRNA-binding sites in the donor template was mutated (TGA) to prevent Cas9-directed cleavage following homology-directed repair. The mStayGold::C4 linker::V5-tag fragment was amplified from pRSETB/mStayGold(c4) (#212018; Addgene) (Ando et al., 2024) with V5-tag harboring primers. DNA fragments were amplified using PCR with the following primers: Bruce 5′ homology arm (forward primer, 5′-TGAAGGTCTCCTTAAAAGCCCGATTGGCGGCGCAGGAAGT-3′, and reverse primer, 5′-TCTTTCTAGGGTTAATCGAGTGCCCAGGGGGCGCGCGAGT-3′), Bruce 3′ homology arm-1 (forward primer, 5′-TCTTTCTAGGGTTAATTTTTGGCATTCACAGGCGATTGCT-3′, and reverse primer, 5′-TCCGGAGGTGCGCCGCGGCAATTTG-3′), mStayGold-C4 linker-V5-tag (forward primer, 5′-CGGCGCACCTCCGGAATGGTGTCTACAGGCGAGGAGCTGT-3′, and reverse primer, 5′-GGTGCTGTCCAGGCCCAGCAGGGGGTTGGGGATGGGCTTGCCCACGGCAGAAGCAGAAGGCTCGTGC-3′), 3′ homology arm-2 with mutated PAM (forward primer, 5′-GGCCTGGACAGCACCGCCACGGAGCAGCATCATCAGCAGCGCGTCCTCAA-3′, and reverse primer, 5′-GACGGCTCTTCATTAACTCTTAAATCAAATTTAAATTGAA-3′). The established plasmids were injected into *yw; nos-Cas9(II-attP40)* for CRISPR/Cas9-mediated transgenesis (BestGene Inc.). Each DsRed-positive transformant was isogenized and confirmed using genomic PCR and sequencing.

All established plasmids were sequenced by Eurofins, Inc. The detailed plasmid DNA sequences are available upon request.

### Cell culture

*Drosophila* S2 cells were grown at 25°C in Schneider’s *Drosophila* medium (GIBCO, 21720001) supplemented with 10% (v/v) heat-inactivated fetal bovine serum, 100 U/mL penicillin, and 100 μg/mL streptomycin (WAKO, 168-23191).

### mNeonGreen signal quantification of S2 cells

*Drosophila* S2 cells (2.5 × 10^5^) seeded in 24-well plates were transfected with 200 ng of the desired plasmid using Effectene Transfection Reagent (QIAGEN, 301427) following the manufacturer’s protocol. Sixteen hours after transfection, cells were transferred to four-compartment glass-bottomed dishes (Greiner #627870). After another 24 hours, images were captured using a Leica TCS SP8 confocal microscope (Leica Microsystems).

The mNeon Green intensity was quantified using Fiji (ImageJ) software (NIH Image). Images were analyzed automatically using the ImageJ macro, as previously described (Fujisawa et al., 2020) with modifications. Briefly, the mNeonGreen channel was filtered by the “Gaussian Blur (sigma = 1)”, processed with “Auto Threshold (“Mean” method),” processed with “Watershed,” and processed with “Analyze particles (size = 40–200 µm^2^, circularity = 0.2–1.0)” to define the region of interest. For a given region of interest, the average mNeonGreen intensity was measured.

### Protein preparation of Drosophila tissues

Whole larvae without digestive tract and fatbodies, larval salivary glands, fatbodies, brains, wing discs and whole adults, adult thoraxes, ovaries, and testes of desired genotype were dissected and lysed with RIPAi buffer (50 mM Tris-HCl, pH 8.0, 150 mM sodium chloride, 0.5 wt/vol% sodium deoxycholate, 0.1% w/v sodium dodecyl sulfate, and 1.0% w/v NP-40) supplemented with cOmplete ULTRA EDTA-free protease inhibitor cocktail (#05892953001, Roche). Samples were homogenized and centrifuged at 20,000 × *g*, 4°C for 10 min. The supernatants were collected and snap-frozen in liquid nitrogen. Protein concentrations were determined using the BCA assay (#297-73101, Wako) following the manufacturer’s protocol. The samples were mixed with Laemmli buffer, boiled at 95°C for 5 min, and then subjected to western blotting analysis.

### Western blotting

Each sample was separated using SDS-PAGE. The proteins were then transferred onto Immobilon-P polyvinylidene fluoride membranes (#IPVH00010; Millipore) for immunoblotting. Membranes were blocked with 4% skim milk diluted in 1× TBST. Immunoblotting was performed using the antibodies mentioned below, which were diluted with 4% skim milk. Signals were visualized using Immobilon Western Chemiluminescent HRP Substrate (#WBKLS0500; Millipore) and FUSION SOLO. 7S. EDGE (Vilber-Lourmat). Contrast and brightness adjustments were applied equally using Fiji (ImageJ) software (NIH Image).

The primary antibodies used included rabbit anti-GFP polyclonal antibody (1:2,000, #598, MBL), mouse anti-V5 tag monoclonal antibody (1:1,000, #46-0705, Invitrogen), mouse anti-FLAG M2 monoclonal antibody (1:5,000, Sigma), rabbit anti-CCT8 polyclonal antibody (1:5,000, this study), rabbit anti-Myc-Tag (71D10) monoclonal antibody (1:5,000, #2278, CST), rabbit anti-3xDYKDDDDK Tag (3xFLAG Tag) (E7C5T) monoclonal antibody (1:5,000, #87537, CST), mouse anti-alpha tubulin (DM1A) monoclonal antibody (1:10,000, #T9026, Sigma), mouse anti-Histone H3 (1B1B2) monoclonal antibody (1:10,000, #14269, CST), mouse anti-Actin monoclonal antibody (1:5,000; #A4700, Sigma), rabbit anti-alpha tubulin polyclonal antibody (1:2,000, #ab15246, Abcam), and rabbit anti-eEF2 polyclonal antibody (1:2,000, #ab33523, Abcam). The secondary antibodies used were goat anti-rabbit IgG horseradish peroxidase (HRP)-conjugated antibody (1:5,000, #7074S, CST) and goat/rabbit/donkey anti-mouse IgG HRP Conjugate (1:5,000, #W402B, Promega). The membranes for streptavidin blotting were blocked with 3% bovine serum albumin. Streptavidin blotting was performed using streptavidin-HRP (1:10,000, #SA10001; Invitrogen) diluted in 3% bovine serum albumin.

### Experimental design for liquid chromatography–mass spectrometry (LC-MS/MS) analyses

Two biological replicates were analyzed for each experimental condition to determine the abundance of Drice/Dcp-1::TurboID relative to the control.

### Purification of biotinylated proteins

Samples were prepared as previously described (Shinoda et al., 2023). Briefly, 100 µg of biotinylated protein-containing lysate was subjected to FG-NeutrAvidin bead (#TAS8848 N1171, Tamagawa Seiki) purification. FG-NeutrAvidin beads (25 µL, approximately 500 µg) were washed three times with RIPA buffer. Benzonase-treated biotinylated protein samples suspended in 1 mL of RIPA buffer were incubated overnight at 4°C. Conjugated beads were magnetically isolated and washed with 500 µL of an ice-cold RIPA buffer solution, 1 M KCl solution, 0.1 M Na_2_CO_3_ solution, and 4 M urea solution. For western blot analyses, the purified samples were mixed with Laemmli buffer, boiled at 95°C for 5 min, and then subjected to western blot analysis. For LC-MS/MS analysis, the purified samples were washed with 500 µL ultrapure water (#214-01301, Wako) and 500 µL of 50 mM ammonium bicarbonate. The samples were then mixed with 50 µL of 0.1% Rapigest diluted in 50 mM ammonium bicarbonate, and 5 µL of 50 mM TCEP was subsequently added. Samples were incubated at 60°C for 5 min, and then 2.5 µL of 200 mM methyl methanethiosulfonate was added. One microgram sequence-grade modified trypsin (#V5111, Promega) was then added for on-bead trypsin digestion at 37°C, leaving the treatment for 16 h. The beads were then magnetically isolated, and 60 µL of the supernatant was collected. Then, 3 µL of 10% trifluoroacetic acid was added to the supernatants, incubating them at 37°C for 60 min with gentle agitation. The samples were then centrifuged at 20,000 × *g*, 4°C for 10 min. The peptides were then desalinated and purified using a GL-tip SDB (#7820-11200, GL Sciences) following the manufacturer’s instructions. The samples were speed-backed at 45°C for 30 min and dissolved in 25 µL of 0.1% formic acid. The samples were then centrifuged at 20,000 × *g*, 4°C for 10 min, and the supernatants were collected. The peptide concentrations were determined using the BCA assay. Finally, 500 ng of the purified protein was subjected to LC-MS/MS analysis.

### LC-MS/MS analyses

Samples were loaded onto an Acclaim PepMap 100 C18 column (75 µm x 2 cm, 3 µm particle size and 100 Å pore size; #164946, Thermo Fisher Scientific) and separated on a nano-capillary C18 column, (75 µm x 12.5 cm, 3 µm particle size, #NTCC-360/75-3-125, Nikkyo Technos) using EASY-nLC 1200 system (Thermo Fisher Scientific). Elution conditions are listed in Table S5. The separated peptides were analyzed using QExactive (Thermo Fisher Scientific) in data-dependent MS/MS mode. The parameters for the MS/MS analysis are listed in Table S6. The collected data were analyzed using Proteome Discoverer (PD) 2.2 software with the Sequest HT search engine. The parameters for the PD 2.2 analysis are described in Table S7. Peptides were filtered at a false discovery rate of 0.01 using the Percolator node. Label-free quantification was performed based on the intensities of the precursor ions using a precursor-ion quantifier node. Normalization was performed using the total amount of peptides in all average scaling modes. MS proteomic data were deposited in the ProteomeXchange Consortium via the jPOST partner repository with the dataset identifier PXD067425 (Okuda et al., 2017).

### Comparison of Dcp-1 interactors

Twenty-four Dcp-1 interactors identified by immune-affinity purification (IAP)-MS (Choutka et al., 2017) were individually searched among all proteins identified in our TurboID-mediated proteomics analysis using their corresponding UniProt accessions. Fifteen proteins were identified in both datasets, and their abundance ratios and #PSMs are listed in Table S2.

### Anti-CCT8 antibody generation

CCT8 was expressed in *Escherichia coli BL21* transformed with *pCold I-EcoRI-CCT8-EcoRI*. Cells were grown at 37 °C overnight, diluted to OD_600_ 0.2 and cultured until OD_600_ 0.5 at 37°C. Protein expression was induced by the addition of 1 mM IPTG at 15°C for 24 h. Cells were centrifuged at 4000 × *g*, 4 °C for 20 min and resuspended in phosphate-buffered saline (PBS). Cells were centrifuged at 4000 × *g*, 4°C for 10 min. Purification of 6x His-tagged CCT8 was performed using the QIAexpress Ni-NTA Fast Start Kit (#30600, Qiagen) according to the manufacturer’s protocol: purification of 6xHis-tagged proteins under denaturing conditions. Briefly, the cell pellets were lysed in Lysis Buffer (Denaturing Buffer adjusted to pH 8.0) with gentle agitation for 60 min. The sample was centrifuged at 9,200 × *g*, 20°C for 30 min. The supernatant was applied to Fast Start column and washed twice with Wash Buffer (Denaturing Buffer adjusted to pH 6.3). Proteins were eluted using Elution Buffer (Denaturing Buffer adjusted to pH 4.5). Polyclonal antibodies were raised in rabbits using purified 6xHis-tagged CCT8 as an immunogen (Eurofins Inc.).

### Protein complexes prediction with AlphaFold3

Amino acid sequences were obtained from the FlyBase database [https://flybase.org/]. The protein complex structures were predicted using AlphaFold3 (Abramson et al., 2024) on the AlphaFold server [https://alphafoldserver.com/].

### Co-immunoprecipitation

*Drosophila* S2 cells (2.0 × 10^6^) seeded in 6-well plates were transfected with the desired plasmid using Effectene Transfection Reagent. For live cell lysate, *Drosophila* S2 cells were lysed on ice using 550 µL of IP lysis buffer (50 mM Tris-HCl (pH 7.5), 150 mM NaCl, 0.5% NP-40) supplemented with cOmplete ULTRA EDTA-free protease inhibitor cocktail. For apoptotic cell lysate, the cells were treated with 10 μg/mL cycloheximide (#C7698, Sigma) for apoptosis induction. After 6 h of incubation, cells were harvested and centrifuged at 800 × *g*, 4 °C for 5 min and washed twice. Then, cells were lysed by using 550 µL of IP lysis buffer (50 mM Tris-HCl (pH 7.5), 150 mM NaCl, and 0.5% NP-40) supplemented with cOmplete ULTRA EDTA-free protease inhibitor cocktail. The cell lysates were centrifuged at 20,000 × *g*, 4 °C for 5 min. Supernatants (50 μL) were collected as inputs. The remainder of the supernatant was incubated for 2 h at 4°C with 20 μL of anti-DDDDK-tag mAb-Magnetic Agarose (#M185-10R, MBL) equilibrated with the IP lysis buffer. The beads were washed thrice in IP lysis buffer and boiled for 5 min at 95°C with 50 μL 1× Laemmli buffer. The samples were magnetically separated, and the supernatants were subjected to SDS-PAGE.

Fifty wing imaginal discs were dissected and lysed on ice using 550 µL of IP lysis buffer supplemented with cOmplete ULTRA EDTA-free protease inhibitor cocktail. Samples were homogenized and centrifuged at 20,000 × *g*, 4°C for 5 min. Supernatants (50 μL) were collected as inputs. The remainder of the supernatant was incubated for 2 h at 4°C with 20 μL of anti-V5-tag mAb-Magnetic Agarose (#M167-10, MBL) equilibrated with the IP lysis buffer. The beads were washed thrice in IP lysis buffer and boiled for 5 min at 95°C with 50 μL 1× Laemmli buffer. The samples were magnetically separated, and the supernatants were subjected to SDS-PAGE.

### Immunohistochemistry

Wing imaginal discs of the late third-instar larvae and testes and ovaries of the adults were fixed with 4% paraformaldehyde in PBS. The wing imaginal discs and testes samples were washed in PBS containing 0.1% Triton X-100 (PBST), and the ovary samples were washed in 1.0% PBST, blocked in PBST with 5% normal donkey serum (PBSTn) for 30 min, and incubated with primary antibodies in PBSTn overnight at 4°C. The samples were washed with PBST, incubated for 2 h with secondary antibodies suspended in PBSTn, and washed again with PBST. The samples were mounted on glass slides with SlowFade Gold antifade reagent (#S36939, Invitrogen). Images were captured using a Leica TCS SP8 microscope (Leica Microsystems). Images were analyzed and edited using the Fiji (ImageJ) software.

The primary antibodies used were rabbit anti-HA (Y11) polyclonal antibody (1:100, #sc-805, Santa Cruz), mouse anti-HA (12CA5) monoclonal antibody (1:200, #ab1424, Abcam), rabbit anti-cleaved Drosophila Dcp-1 (Asp215) polyclonal antibody (1:200, #9578S, CST), rabbit anti-cleaved PARP (Asp214) (D64E10) monoclonal antibody (1:200, #5625S, CST), mouse anti-Myc-tag monoclonal antibody (1:500, #R950-25, Invitrogen) and mouse anti-V5 tag monoclonal antibody (1:100, #960-25, Invitrogen). The secondary antibodies used were Cy3-conjugated donkey anti-rabbit IgG antibody (1:250, #711-165-152, Jackson Immuno Research), Alexa647-conjugated donkey anti-mouse IgG antibody (1:100, #A-31571, Thermo Fisher Scientific), Alexa488-conjugated donkey anti-rabbit IgG antibody (1:200, #A-21206, Thermo Fisher Scientific), Alexa647-conjucated donkey anti-rabbit IgG antibody (1:200, #A-31573, Thermo Fisher Scientific), Alexa488-conjugated donkey anti-mouse IgG (1:500 (Myc) or 1:100 (V5), #A-21202, Thermo Fisher Scientific). Hoechst 33342 (8 µM; #H3570, Invitrogen) for nuclear staining and Phalloidin-iFluor633 (1:1000, #ab176758, Abcam) for F-actin staining were used during the secondary antibody incubation.

### Terminal deoxynucleotidyl transferase dUTP nick end labeling (TUNEL) assay

TUNEL assay was performed using the ApopTag Red in situ apoptosis detection kit (#S7165, Millipore) according to the manufacturer’s instruction as previously described (Shinoda et al., 2019). Wing imaginal discs were dissected and fixed for 20 min at room temperature in 1% PFA in PBS. The samples were then washed with PBS three times, 5 min per wash. Next, the samples were re-fixed in ethanol:acetic acid, 2:1, for 5 min at -20 °C; washed in PBS twice, 5 min per wash; incubated in Equilibration Buffer for 10 sec; and incubated again in Working Strength TdT Enzyme at 37℃ for 1 h. The TdT reaction mix was replaced with Working Strength Stop/Wash Buffer and incubated for 10 min at room temperature. Then, the samples were washed three times with PBS, 1 min per wash; and incubated with Working Strength Rhodamine Antibody Solution with 8 µM Hoechst 33342 for 4 h at room temperature. The samples were then washed four times in PBS, 2 min per wash. The samples were mounted on glass slides with SlowFade Gold antifade reagent Images were captured using a Leica TCS SP8 microscope. Images were analyzed and edited using the Fiji (ImageJ) software.

### Wing size measurement

Flies were raised, and their wings were mounted and digitized as previously described (Shinoda et al., 2019). Briefly, flies were placed on an acetic acid agar plate (2% agar, 1% sucrose, and 0.36% acetic acid) to lay eggs overnight. The plates were collected, and the eggs were incubated at 25°C for 24 h. The 1st instar larvae were collected from the agar plate, and 30 larvae were placed in one vial to precisely control the larval density. These flies were raised at 25°C. After more than two days of hatching, the adult female flies were collected and submerged in 99.5% ethanol to dehydrate for 24 hours. Dehydrated flies were dissected to mount their wings using Euparal (Waldeck) with the dorsal side upward. Images were digitized using an upright microscope (Leica DM5000B; Leica Microsystems) equipped with a CCD camera. The wing size was measured using a Drosophila wing digital twin [https://datamarkin.com/models/drosophila-wings-beta] developed by Datamarkin. Areas 0, 1, and 2 were quantified as the anterior parts, whereas areas 4, 6, 7, and 8 were quantified as the posterior parts, respectively.

### Statistical analysis

The data are presented as the means ± standard error of measurement (SEM). *p*-values were calculated using a one-way analysis of variance (ANOVA), followed by Sidak’s multiple comparison test for selected pairs (Figure 1–figure supplement 2B, C, D; Figure 5K), or Dunnett’s multiple comparison test (Figure 5G, I; Figure 5–figure supplement 1A) using GraphPad Prism 8. NS: *p* ≧ 0.05, †: *p* < 0.05.

## Acknowledgments

We would like to thank G. W. Davis, R. Carthew, B. Hay, C-H. Cheng, H. Richardson, M. Suzanne, D. W. Williams, S. K. Yoo, H. Steller, Y. Hiromi, N. Fujita, H. D. Ryoo, the FlyORF Zurich ORFeome Project, Bloomington *Drosophila* Stock Center, Vienna Drosophila Resource Center, Kyoto Stock Center, and NIG-FLY for providing the fly strains. We thank Datamarkin for providing the Drosophila wing digital twin tool. We thank Dr. S. Hirayama, Dr. S. Murata, and the One-Stop Sharing Faculty Center for Future Drug Discovery at the Graduate School of Pharmaceutical Sciences, University of Tokyo, for the LC-MS/MS analysis. We thank Miura’s lab members for their technical assistance and discussions; in particular, K. Takenaga for preparing the fly food and R. Takamoto for experimental support. We thank *Editage* [www.editage.com] for the English language editing of this manuscript. This work was supported by grants from the Ministry of Education, Culture, Sports, Science, and Technology of Japan (KAKENHI Grant Numbers 19K16137, 21K15080, 22H05586, and 23K05747 to N.S. and 21H04774, 23H04766, 24H00567, and 25H01842 to M.M.), the Japan Science and Technology Agency (ACT-X, Grant Number JPMJAX2539), the Japan Agency for Medical Research and Development (AMED; Grant number JP21gm5010001 to M.M.), and grants from the Takeda Science Foundation and Sumitomo Foundation to N.S.

## Author Contributions

N.S. and M.M. designed the study; N.S. conducted most of the experiments and analyzed most of the data; Y.H conducted protein structural predictions; N.H. analyzed the expression patterns of Drosophila caspases; and N.S. and M.M. wrote the manuscript.

---

**Figure 1–figure supplement 1.**
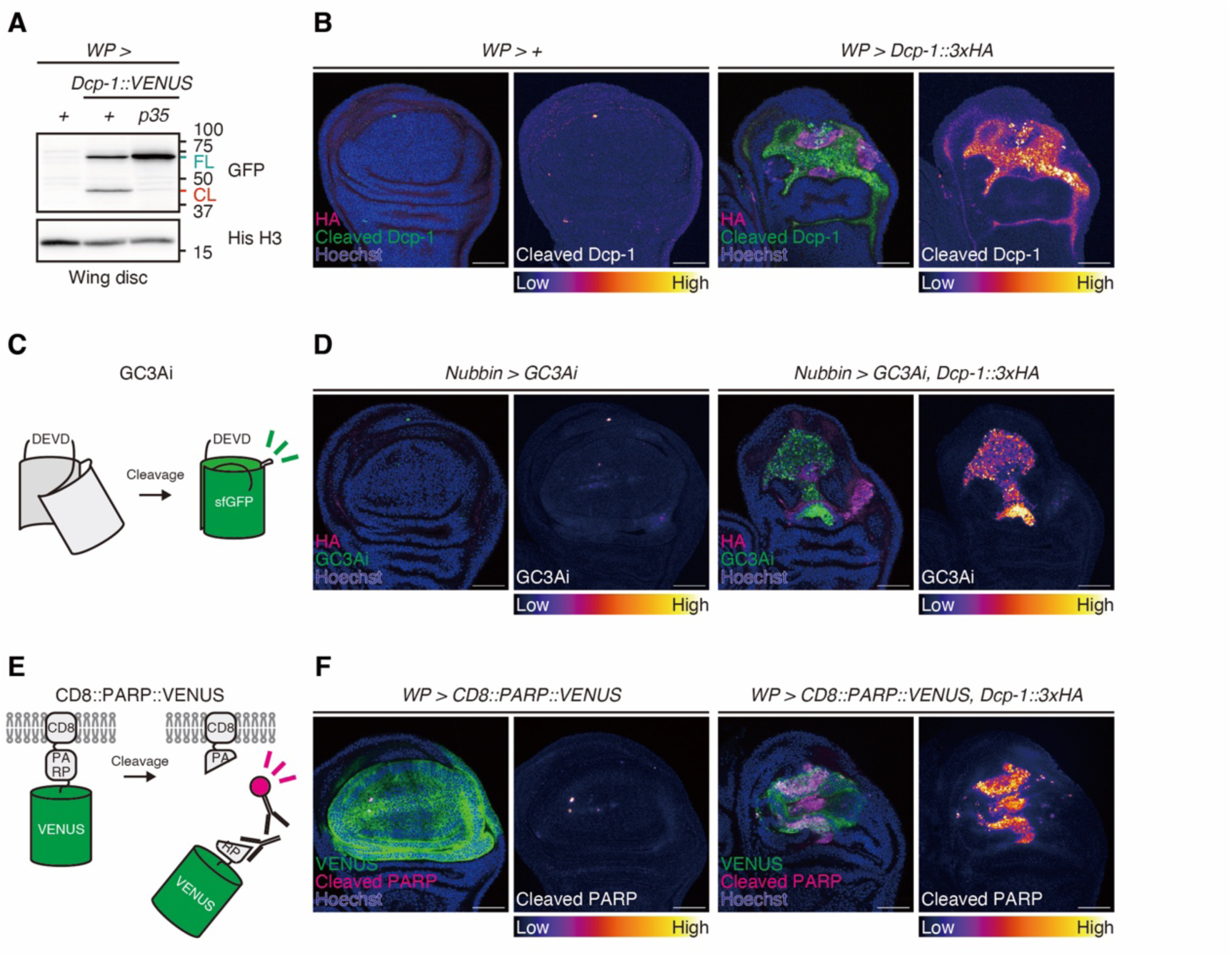
*Dcp-1* overexpression leads to its activation and the cleavage of executioner caspase substrates. (A) Western blotting of the cleavage of C-terminally VENUS tagged Dcp-1 overexpressed in larval wing imaginal disc using *WP-Gal4*. Both full-length (FL) Dcp-1::VENUS (MW: 62.7 kDa) and cleaved (CL) Dcp-1::VENUS (MW: 39.1 kDa) are detected using anti-GFP antibody. (B) Representative images of the anti-cleaved Dcp-1 antibody staining (green) in larval wing imaginal disc upon *3xHA-tagged Dcp-1* (magenta) overexpression using *WP-Gal4*. Nuclei are visualized by Hoechst 33342 (blue). Scale bar: 50 µm. (C) Schematic structures of fluorescent switch-on probe, GC3Ai. Caspase-mediated cleavage makes the probe fluorescent. Caspase activity is measured by the fluorescent intensity of the probe. (D) Representative images of the GC3Ai fluorescent (green) in larval wing imaginal disc upon *3xHA-tagged Dcp-1* (magenta) overexpression using *Nubbin-Gal4*. Nuclei are visualized by Hoechst 33342 (blue). Scale bar: 50 µm. (E) Schematic structures of CD8::PARP::VENUS. Caspase-mediated cleavage generates neo-epitope of anti-cleaved PARP antibody. Caspase activity is measured by the intensity of the immunohistochemistry. (F) Representative images of the anti-cleaved PARP antibody staining (magenta) in larval wing imaginal disc expressing CD8::PARP::VENUS probe (VENUS, green) and *3xHA-tagged Dcp-1* overexpression using *WP-Gal4*. Nuclei are visualized by Hoechst 33342 (blue). Scale bar: 50 µm.

**Figure 1–figure supplement 2.**
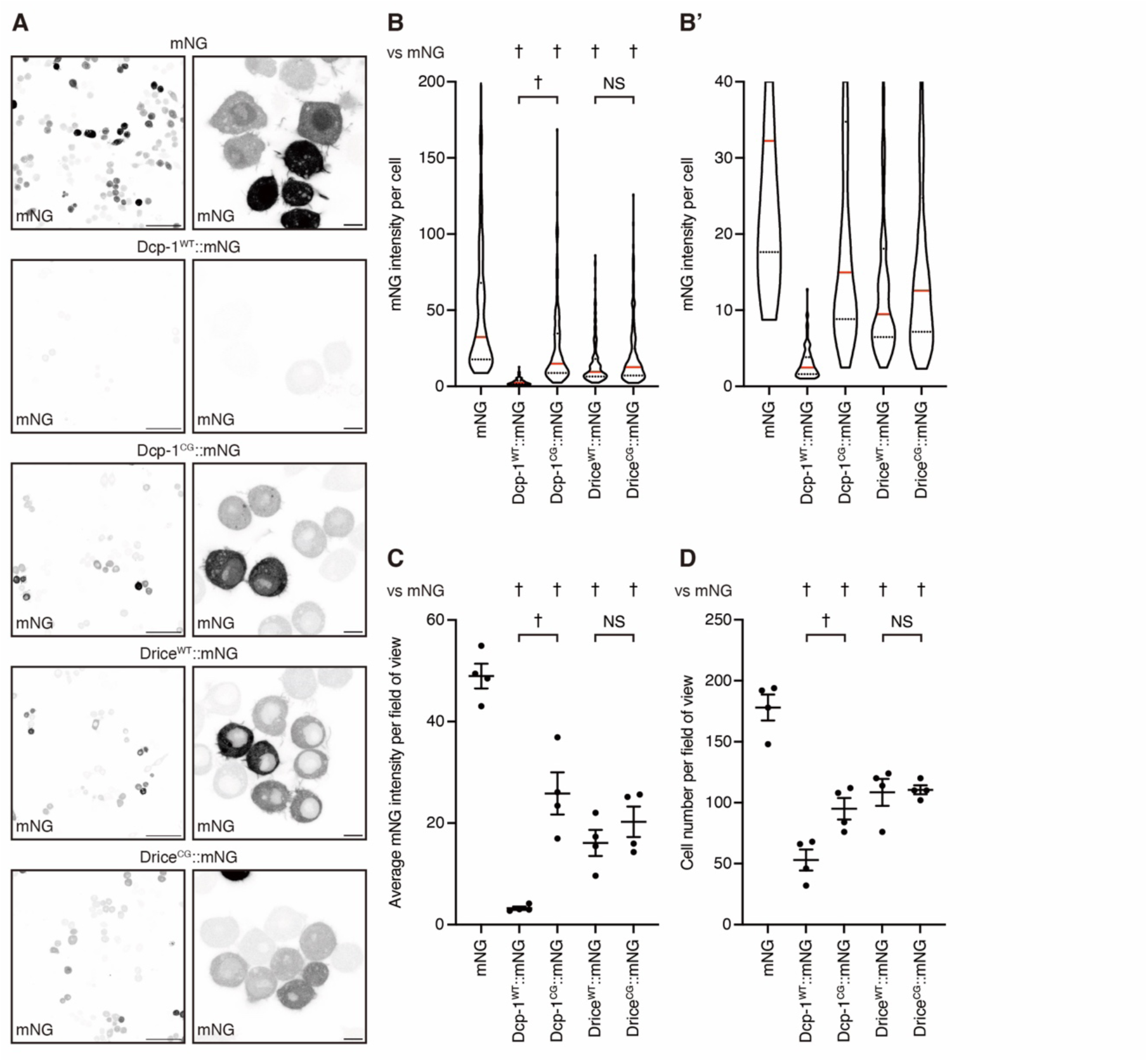
Dcp-1 overexpression induces apoptosis in *Drosophila* S2 cells. (A) Representative images of *Drosophila* S2 cells transfected with mNeonGreen (mNG)-tagged wild-type (WT) or catalytically inactive (catalytic cysteine mutated to glycine; CG) Dcp-1 and Drice. Scale bars: 50 µm (left) and 5 µm (right). (B) Quantification of mNG intensity for each cell, shown as a violin plot. Quartiles are indicated by black broken lines, and the median is indicated by a red bar. (B′) Magnified view of the data in (B), with the Y-axis limited to 0–40. *p*-values were calculated using one-way analysis of variance (ANOVA) followed by Sidak’s multiple-comparison test for selected pairs. NS, *p* > 0.05; †, *p* < 0.05. Sample sizes: mNG (n = 712 cells), Dcp-1^WT^::mNG (n = 212 cells), Dcp-1^CG^::mNG (n = 380 cells), Drice^WT^::mNG (n = 434 cells), and Drice^CG^::mNG (n = 442 cells). (C) Quantification of the average mNG intensity in each field of view. Data are presented as mean ± SEM. *p*-values were calculated using one-way ANOVA followed by Sidak’s multiple-comparison test for selected pairs. NS, *p* ≧ 0.05; †, *p* < 0.05. Sample size for each condition is 4. (D) Quantification of the mNG-positive cell number in each field of view. Data are presented as mean ± SEM. *p*-values were calculated using one-way ANOVA followed by Sidak’s multiple-comparison test for selected pairs. NS, *p* ≧ 0.05; †, *p* < 0.05. Sample size for each condition is 4.

**Figure 2–figure supplement 1.**
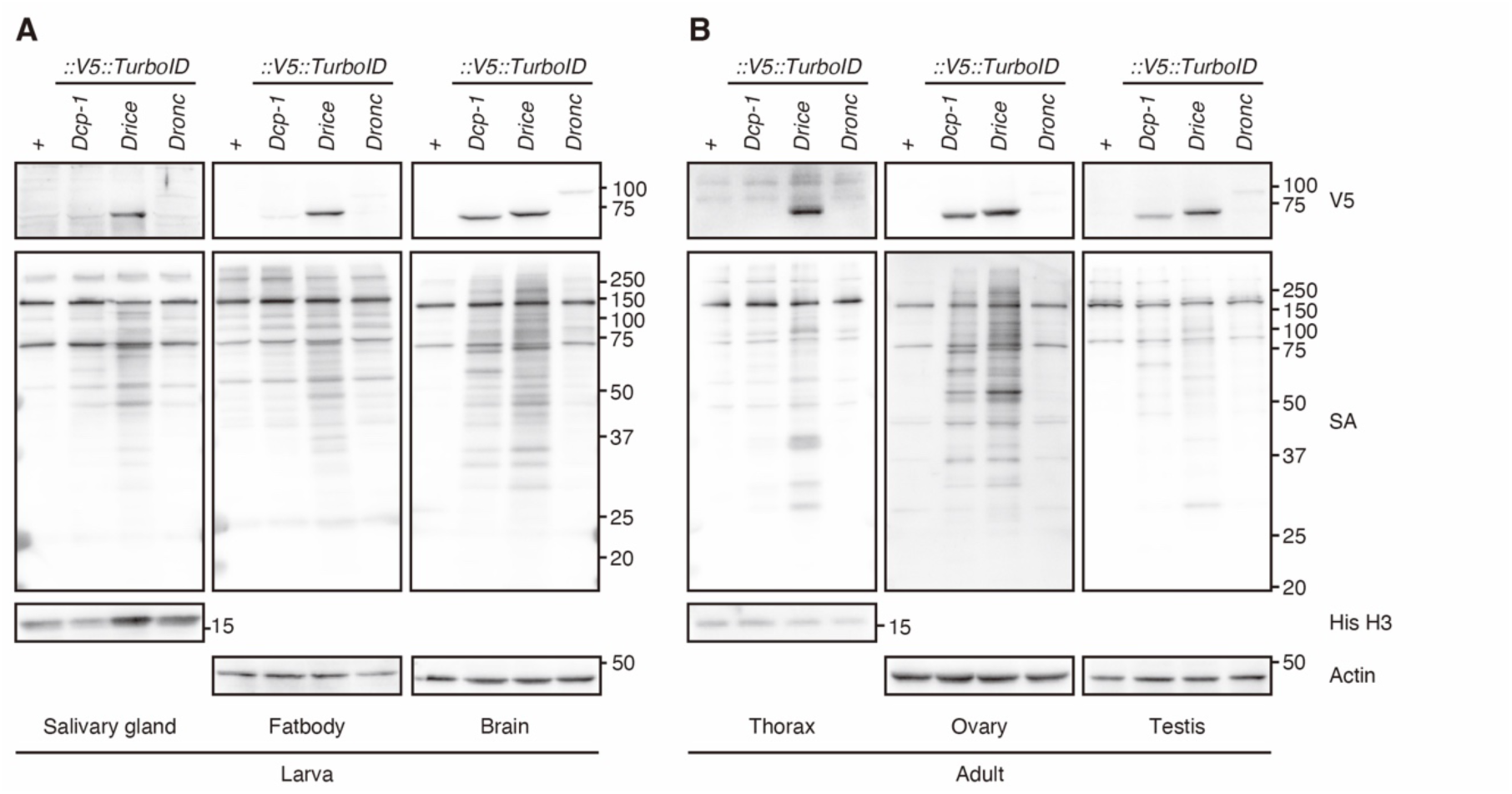
Expression patterns of *Drosophila* caspases in larval and adult tissues. (A) Western blotting of expression of C-terminally V5::TurboID knocked-in tagged caspases and biotinylation patterns of their proximal proteins detected by streptavidin (SA) in larval salivary glands, fat bodies, and brains. (B) Western blotting of expression of C-terminally V5::TurboID knocked-in tagged caspases and biotinylation patterns of their proximal proteins detected by SA in adult thoraxes, ovaries, and testes.

**Figure 2–figure supplement 2.**
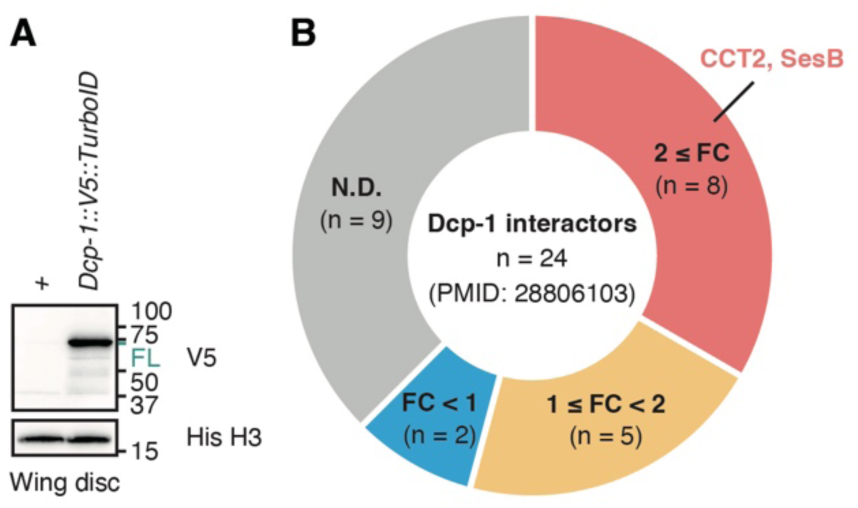
Comparison of Dcp-1 interactors identified by TurboID-MS and immune-affinity purification (IAP)-MS. (A) Western blotting of C-terminally V5::TurboID knocked-in tagged Dcp-1 in larval wing imaginal disc. Only full-length (FL) Dcp-1::V5::TurboID (MW: 72.8 kDa), but not cleaved Dcp-1::V5::TurboID (MW: 49.2 kDa), is detected using anti-V5 antibody. (B) Pie chart of Dcp-1 interactors identified by immunoaffinity purification (IAP)-MS (Choutka et al., 2017). Among the 24 Dcp-1 interactors, 15 were also identified by our TurboID-MS analysis. Proteins are grouped according to their fold change relative to the control (FC [Dcp-1::V5::TurboID/w^1118^]).

**Figure 2–figure supplement 3.**
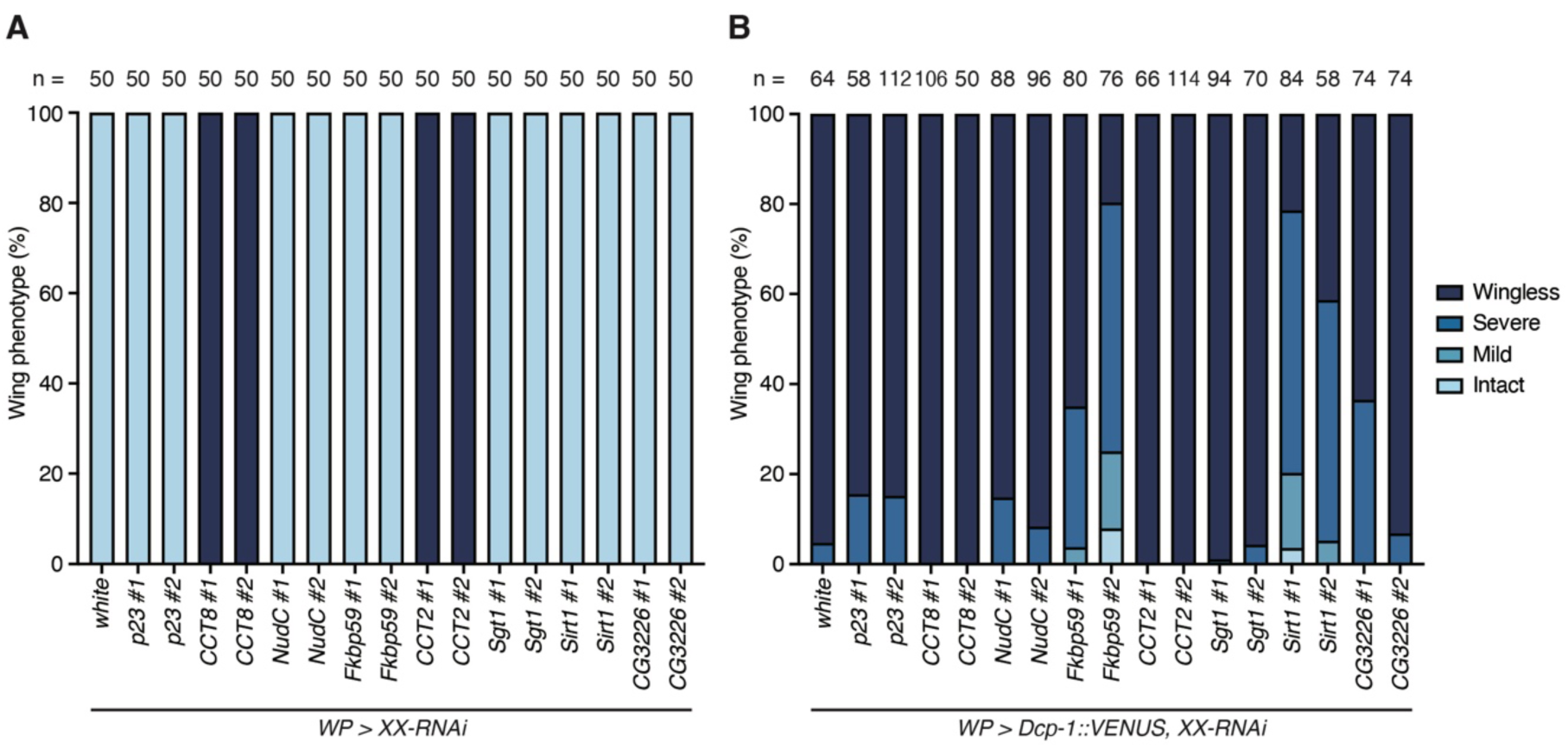
RNAi screening of Dcp-1-enriched proximal proteins for regulators of Dcp-1 activation. (A) Quantification of female wing phenotypes upon knockdown of Dcp-1-enriched proximal proteins driven by *WP-Gal4*. “XX” denotes the knocked-down gene, and “#1” and “#2” indicate independent RNAi lines. Each wing is manually classified into four (wingless, severe, mild, and intact) categories. Sample sizes are shown in the figure. (B) Quantification of female wing phenotypes upon *Dcp-1::VENUS* overexpression driven by *WP-Gal4* with simultaneous knockdown of Dcp-1-enriched proximal proteins. “XX” denotes the knocked-down gene, and “#1” and “#2” indicate independent RNAi lines. Each wing is manually classified into four (wingless, severe, mild, and intact) categories. Sample sizes are shown in the figure.

**Figure 3–figure supplement 1.**
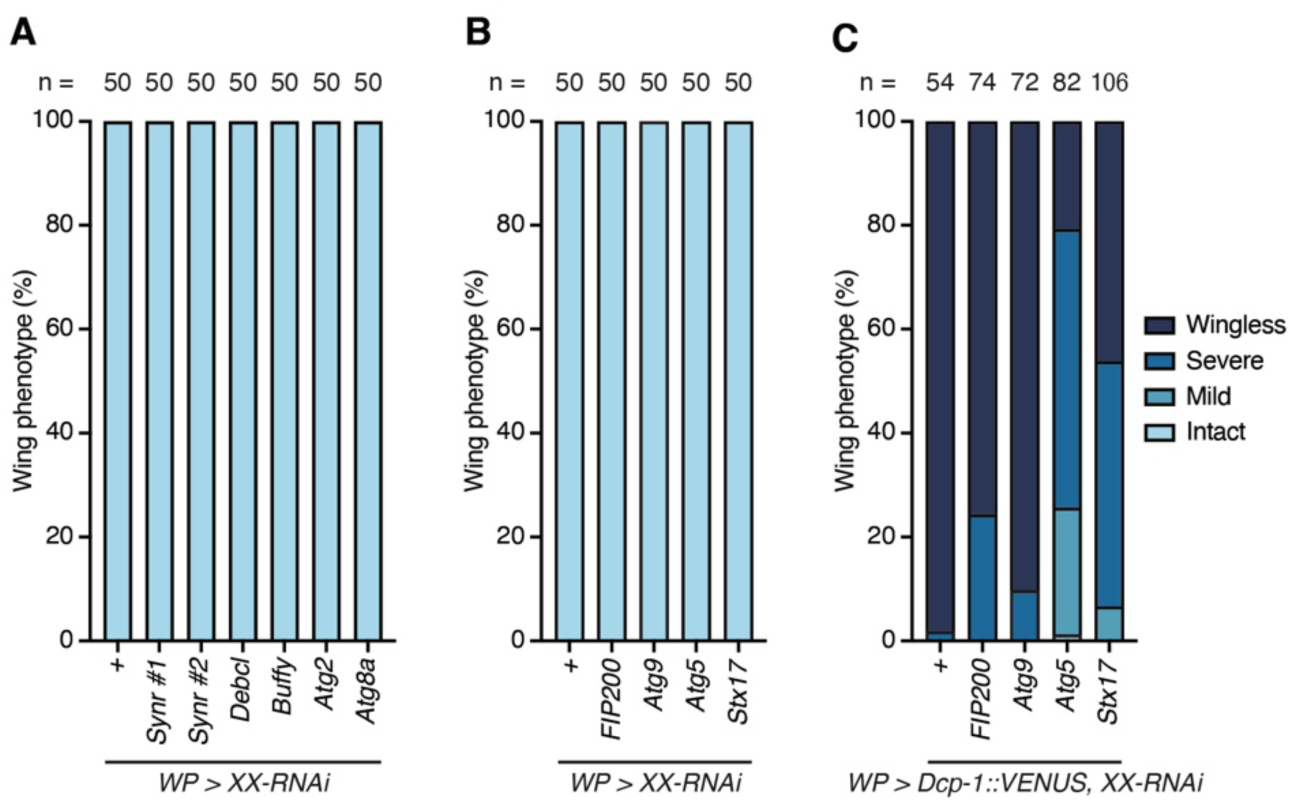
Knockdown of autophagy-related genes for regulators of Dcp-1 activation. (A) Quantification of female wing phenotypes upon knockdown of *Synr*, *Debcl*, *Buffy,* and autophagy-related genes by *WP-Gal4*. “XX” denotes the knocked-down gene. Each wing is manually classified into four (wingless, severe, mild, and intact) categories. Sample sizes are shown in the figure. (B) Quantification of female wing phenotypes upon knockdown of autophagy-related genes by *WP-Gal4*. “XX” denotes the knocked-down gene. Each wing is manually classified into four (wingless, severe, mild, and intact) categories. Sample sizes are shown in the figure. (C) Quantification of female wing phenotypes upon *Dcp-1::VENUS* overexpression driven by *WP-Gal4* with simultaneous knockdown of autophagy-related genes by. “XX” denotes the knocked-down gene. Each wing is manually classified into four (wingless, severe, mild, and intact) categories. Sample sizes are shown in the figure.

**Figure 4–figure supplement 1.**
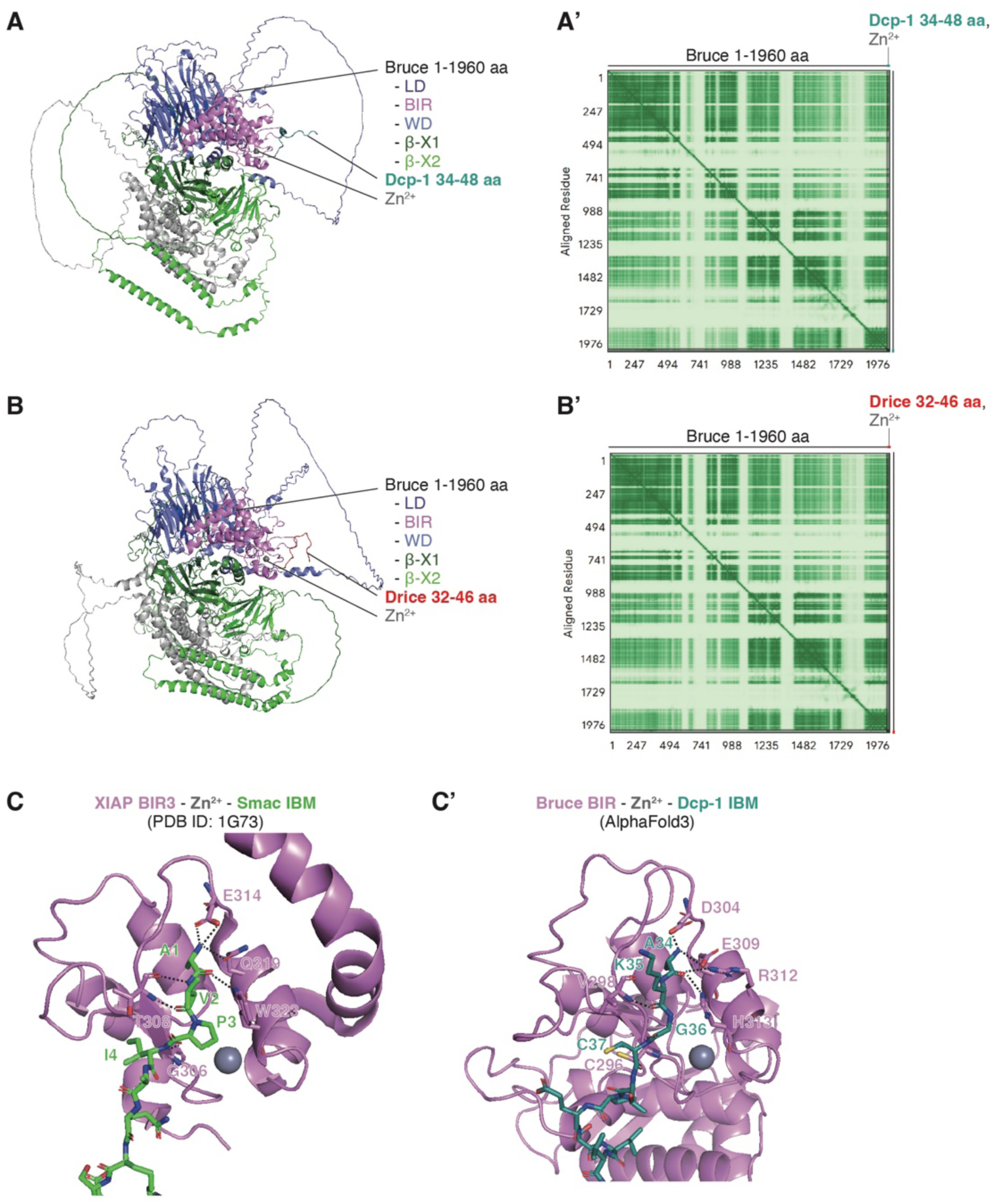
Predicted structures of Bruce complexed with Dcp-1 and Drice IBM-like sequences. (A) Overall view of the predicted complex structure of Bruce and Dcp-1. Model 0 is shown with the same color scheme as in Figure 4A and B. The predicted position error is shown in (A’). (B) Overall view of the predicted complex structure of Bruce and Drice. Model 0 is shown with the same color scheme as in Figure 4A and C. The predicted position error is shown in (B’). (C) Structural comparison between the experimentally determined complex of XIAP BIR3 and Smac IBM (C), and the predicted complex of Bruce BIR and Dcp-1 IBM (C’). XIAP BIR3 is shown as a pink cartoon and the Smac IBM as green sticks (PDB ID: 1G73, right panel). Bruce BIR domain is shown as a pink cartoon and the Dcp-1 IBM as dark green sticks (model 0, left panel). Zn²⁺ ions are shown as gray spheres. Salt bridges are indicated by black dashed lines, along with the corresponding interacting residues.

**Figure 4–figure supplement 2.**
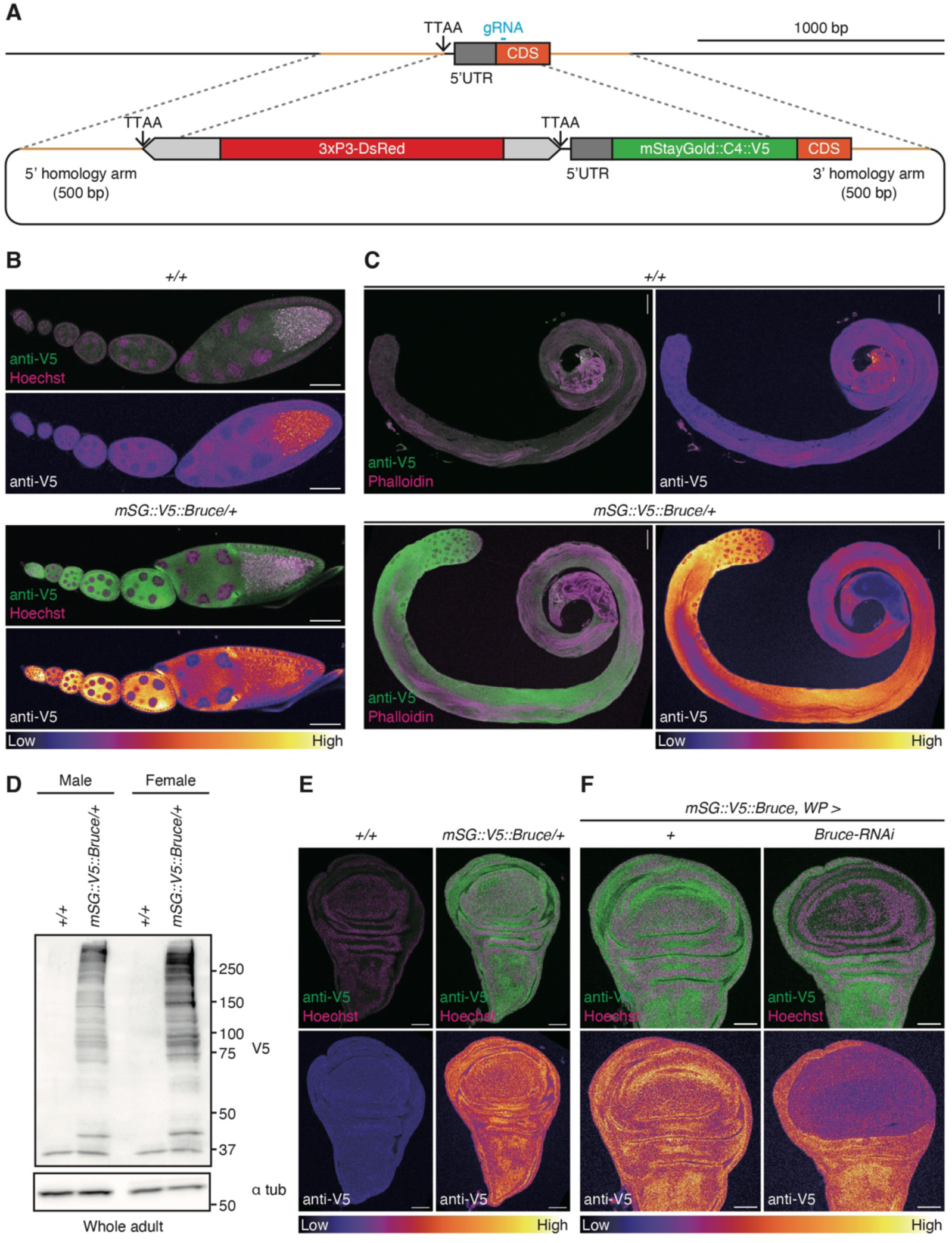
Expression patterns of mStayGold::V5-tag knocked-in Bruce. (A) Schematic diagram of CRISPR/Cas9-mediated knock-in of mStayGold::C4 linker (C4)::V5 tag at the N-terminus of Bruce. A 3xP3-DsRed cassette was integrated at the endogenous TTAA site as a transformation marker. Removal of the cassette restores the genomic TTAA site, enabling scarless genome engineering. (B) Expression pattern of mStayGold (mSG)::C4::V5-tag knocked-in Bruce detected by anti-V5 tag antibody (green) in the ovariole. Nuclei are visualized using Hoechst 33342 (magenta). Scale bar: 50 µm. (C) Expression pattern of mSG::C4::V5-tag knocked-in Bruce detected by anti-V5 tag antibody (green) in the testis. F-actin is visualized using Phalloidin (magenta). Scale bar: 50 µm. (D) Western blotting of expression of mSG::C4::V5-tag knocked-in tagged Bruce in the whole adult male and female. (E) Expression pattern of mSG::C4::V5-tag knocked-in tagged Bruce detected by anti-V5 tag antibody (green) in the wing imaginal discs. Nuclei are visualized using Hoechst 33342 (magenta). Scale bar: 50 µm. (F) Expression pattern of mSG::C4::V5-tag knocked-in tagged Bruce detected by anti-V5 tag antibody (green) in the wing imaginal discs upon *Bruce-RNAi* using *WP-Gal4*. Nuclei are visualized using Hoechst 33342 (magenta). Scale bar: 50 µm.

**Figure 5–figure supplement 1.**
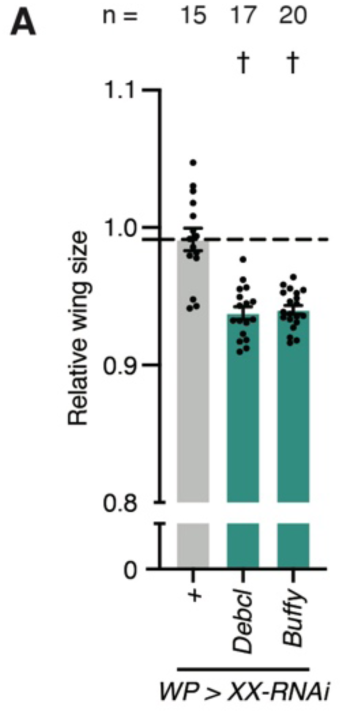
Effect of *Debcl* or *Buffy* knockdown on wing tissue growth. (A) Quantification of relative wing size normalized to the corresponding *No-Gal4* control. Data are presented as mean ± SEM. *p*-values were calculated using one-way analysis of variance (ANOVA) followed by Dunnett’s multiple-comparison test against no UAS control. †, *p* < 0.05. Sample sizes are shown in the figure.

## Legends for Supplemental Tables

**Table S1. Dcp-1 and Drice proximal protein lists.**

Lists of the identified Dcp-1 proximal proteins (abundance ratio (Dcp-1::V5::TurboID/Drice::V5::TurboID) > 2), Drice proximal proteins (abundance ratio (Drice::V5::TurboID/Dcp-1::V5::TurboID) > 2), and candidates for RNAi screening (abundance ratio (Dcp-1::V5::TurboID/Drice::V5::TurboID) > 1.5, abundance ratio (Dcp-1::V5::TurboID/Control) > 10, PSM ≧ 2). The UniProt accessions, gene descriptions, abundance ratios, and # PSMs are listed.

**Table S2. Comparison of Dcp-1 interactors.**

List of previously reported Dcp-1 interactors and comparison with Dcp-1 proximal proteins identified by TurboID-MS. The UniProt accessions, gene descriptions, abundance ratios, and # PSMs are listed.

**Table S3. Fly stock list.**

List of fly strains. Genotypes, maps, source or references, and stock numbers are listed.

**Table S4. Detailed genotypes.**

Detailed genotypes and their related figure # are presented.

**Table S5. Liquid chromatography (LC) settings.**

Gradient settings for LC analysis are presented.

**Table S6. Tandem mass spectrometry (MS/MS) settings.**

Settings used for the MS/MS analysis are presented.

**Table S7. Proteome Discoverer 2.2 settings.**

Proteome Discoverer 2.2 settings for proteomics analysis are presented.

